# Spinal V2b neurons reveal a role for ipsilateral inhibition in speed control

**DOI:** 10.1101/615906

**Authors:** Rebecca A. Callahan, Richard Roberts, Mohini Sengupta, Yukiko Kimura, Shin-ichi Higashijima, Martha W. Bagnall

**Affiliations:** Washington University School of Medicine, Department of Neuroscience, St. Louis, MO, USA; National Institute for Basic Biology, Okazaki, Aichi, Japan

## Abstract

The spinal cord contains a diverse array of interneurons that govern motor output. Traditionally, models of spinal circuits have emphasized the role of inhibition in enforcing reciprocal alternation between left and right sides or flexors and extensors. However, recent work has shown that inhibition also increases coincident with excitation during contraction. Here, using larval zebrafish, we investigate the V2b (Gata3+) class of neurons, which contribute to flexor-extensor alternation but are otherwise poorly understood. Using newly generated transgenic lines we define two stable subclasses with distinct neurotransmitter and morphological properties. These two V2b subclasses make direct synapses onto motor neurons with differential targeting to slower and faster circuits. *In vivo*, optogenetic suppression of V2b activity leads to increases in locomotor speed. We conclude that V2b neurons exert speed-specific influence over axial motor circuits throughout the rostrocaudal axis. Together, these results indicate a new role for ipsilateral inhibition in speed control.

## Introduction

Rhythmic, coordinated body movements require selective recruitment of motor neurons by spinal and supraspinal premotor circuits. Most vertebrates locomote via alternating left-right contractions that travel from rostral to caudal; tetrapods additionally alternate between flexors and extensors to regulate limb movements. Due in part to the technical challenges in identifying and manipulating specific classes of neurons in the spinal cord, the underlying circuitry of locomotion remains only poorly worked out.

Spinal premotor neurons are broadly divided into five superclasses arising from distinct progenitor domains (dI6, V0, V1, V2, V3) [1]. Within these superclasses, cardinal neuron classes have been identified based on transcription factor expression and neurotransmitter identity (e.g., V2a / Chx10 / excitatory; V2b / Gata3 / inhibitory). Recently, it has become clear that many of these classes can be further subdivided into anywhere from 2 to 50 subclasses, based on anatomical and genetic distinctions, with as-yet unclear implications for circuit connectivity and function [2–6].

Traditionally, patterned locomotion has been modeled as an alternation between excitation and inhibition, which dominate motor neurons during contraction and extension portions of the cycle, respectively [7, 8]. Recently, however, evidence from both fish and turtles has challenged the notion that inhibition is minimal during the contraction of the cycle, i.e. in-phase with excitation. Instead, inhibitory conductances appear to be significant both in- and out-of-phase [9–13], suggesting that simultaneous recruitment of excitation and inhibition during the contraction is important for regulating motor neuron firing [14].

In-phase inhibition is thought to derive from two spinal interneuron classes, the V1 and V2b populations. The V1 population includes Renshaw cells [15, 16], which provide recurrent inhibition onto motor neurons with potentially significant shunting effects [17]. To date, most analysis of drive from V2b neurons has focused on the shared contributions of V1s and V2bs to reciprocal inhibition governing flexor/extensor alternation in limbed animals [18–20]. However, this does not shed light on potential functions of ipsilateral inhibition in gain control for regulation of motor neuron firing *during* contraction, as opposed to suppression of motor neuron firing during extension.

In-phase inhibition increases in amplitude for faster locomotor movements [10] suggesting a potential role in speed control. Here we investigated whether V2b neurons could indeed provide direct inhibition to motor neurons for speed control, taking advantage of the speed-dependent organization of zebrafish motor circuits [21–23]. V2b neurons are good candidates for in-phase gain control because they are exclusively inhibitory in mouse and zebrafish [24] with ipsilateral, descending axons within the spinal cord [18, 25]. They arise from a final progenitor division that produces pairs of V2a and V2b neurons [26]. Given the role of V2a neurons in triggering motor output [27, 28], particularly through speed-specific circuits for titrating levels of motor excitation [4, 29-31], it seems plausible that their sister V2b neurons exert an opposing, inhibitory role in speed control. However, the V2b class has not been well characterized at anatomical or neurochemical levels outside of very early development.

Here, we define two subclasses of V2b neurons in larval zebrafish based on differential transmitter expression and anatomy, and further show that these neurons directly inhibit axial motor neurons in speed-specific circuits. Optogenetic suppression of V2b activity elicits faster locomotion, consistent with a new role for ipsilateral inhibition in speed control.

## Results

### Gata3 transgenic lines label V2b neurons

V2b neural identity is, in part, conferred by the developmental expression of the transcription factor Gata3 [19, 32]. To provide transgenic labeling of the V2b population, we generated two *gata3* transgenic lines, *Tg(gata3:loxP-DsRed-loxP:GFP)* and *Tg(gata3:Gal4)* from bacterial artificial chromosomes (BAC) insertion transgenesis. Both lines label V2b neurons throughout the rostrocaudal extent of the larval zebrafish spinal cord (Fig. 1A; *Tg(gata3:loxP-DsRed-loxP:GFP)* line shown). *Gata3*-driven fluorescent proteins are also broadly expressed in the brain, hindbrain, and assorted non-nervous system soft tissue including the pronephric duct [33].

**Figure 1.**
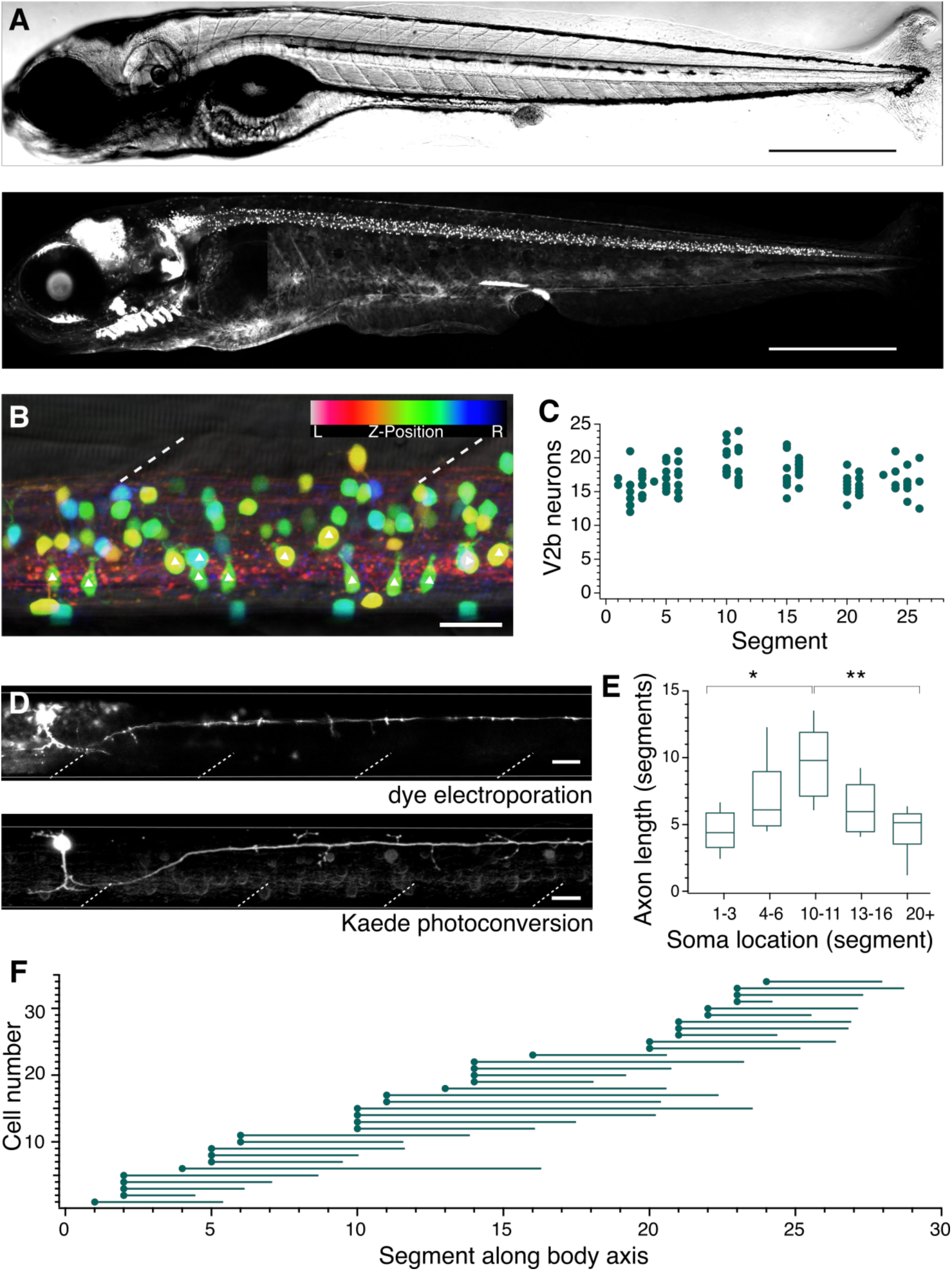
V2b neurons are found throughout the rostral-caudal axis of zebrafish spinal cord. (A) Transmitted DIC image (top) and confocal image (bottom) of a 5dpf *Tg(gata3:loxP-dsRed-loxP:GFP)* animal. Scale bars = 0.5 mm. (B) Lateral view of a midbody spinal cord segment, false color depth-coded from left to right; dashed lines mark muscle segments. In this and all subsequent figures, rostral is to the left and dorsal is to the top. Triangles mark CSF-cN neurons. Scale bar = 20 μm. (C) V2b cell counts per hemisegment quantified along the rostrocaudal body axis, n = 7 fish. (D) Example cell morphology using two techniques to label single V2b axons: single-cell dye electroporation (top) and Kaede photoconversion (bottom). Scale bar = 20 μm. (E) Midbody V2b neurons extend axons through more segments than V2b neurons in other rostrocaudal locations. *p < 0.01; **p < 0.001, ANOVA and Tukey’s test. (F) Ball and stick plots indicate soma position and axon extension along the body axis for 35 V2b neurons.

In a typical spinal segment, V2b soma position ranged from medial to lateral, as visualized with a color depth code (Fig. 1B). Gata3 is expressed in not only V2b neurons but also the mechanosensory cerebrospinal fluid contacting neurons (CSF-cN)[34]. CSF-cNs have a distinct anatomy including large soma size, ventral position, and stereotyped extension into the central canal (triangles, Fig. 1B), permitting straightforward exclusion from further V2b analysis. On average, each hemisegment contained 17.2 +/- 2.5 (mean +/- SD) V2b neuron somata with relatively little variation from rostral to caudal segments (Fig. 1C).

### V2b axons extend throughout the spinal cord

In V2b axons To visualize V2b axonal trajectories within the spinal cord, we labeled individual neurons via either single cell dye-electroporation or Kaede photoconversion in a *Tg(gata3:Gal4, UAS:Kaede)* line (Fig. 1D) [35]. No difference in axon length or trajectory was observed between the two methods. In all 59 neurons, the axon descended caudally and ipsilaterally, with an extent ranging from 2 – 15 segments. V2b axons originated on the ventral aspect of the soma and projected laterally into the white matter. Putative en passant boutons were seen as swellings distributed along the axon. Most V2b axons projected short collaterals into the soma-dense medial spinal cord along the axon extent. V2b dendrites extended from the main axon branch near the soma, (Fig. 1D), similar to identified mixed processes in V2a neurons [4]. However, in contrast to V2a neurons[4], no V2b neurons extended rostral axons beyond the segment of origin.

Single-cell Kaede photoconversions made at different positions along the rostrocaudal extent of the spinal cord revealed that axonal projections were longest for V2b somata located in the midbody range (Figs. 1E and 1F). Overall, these data reveal that zebrafish V2b neurons exclusively innervate areas ipsilateral and caudal to the soma, with the greatest territory of axonal coverage originating from mid-body neurons with long axons.

### In situ hybridization validates transgenic animal lines

To validate the transgenic lines used in this work, we performed two-color fluorescent in-situ hybridization on each line, examining whether the fluorescent reporter expression matched with RNA expression of the targeted gene. We evaluated *completeness* of label, i.e. the percentage of neurons expressing the endogenous gene that also express the fluorescent reporter, and *accuracy* of label, i.e. the percentage of fluorescent reporter-expressing neurons that express the targeted endogenous gene. These metrics were evaluated for *Tg(gata3:loxP-DsRed-loxP:GFP)*, *Tg(gata3:Gal4,UAS:GFP)*, *Tg(gad1b:GFP)*, and *Tg(glyt2:loxP-mCherry-loxP:GFP)*, n = 4-6 animals for each line. An example of each is provided in Figure 2. Results for the completeness and accuracy of transgenic lines are reported in Table 1. All lines were sufficiently complete and accurate for use in further quantitative analyses.

**Figure 2.**
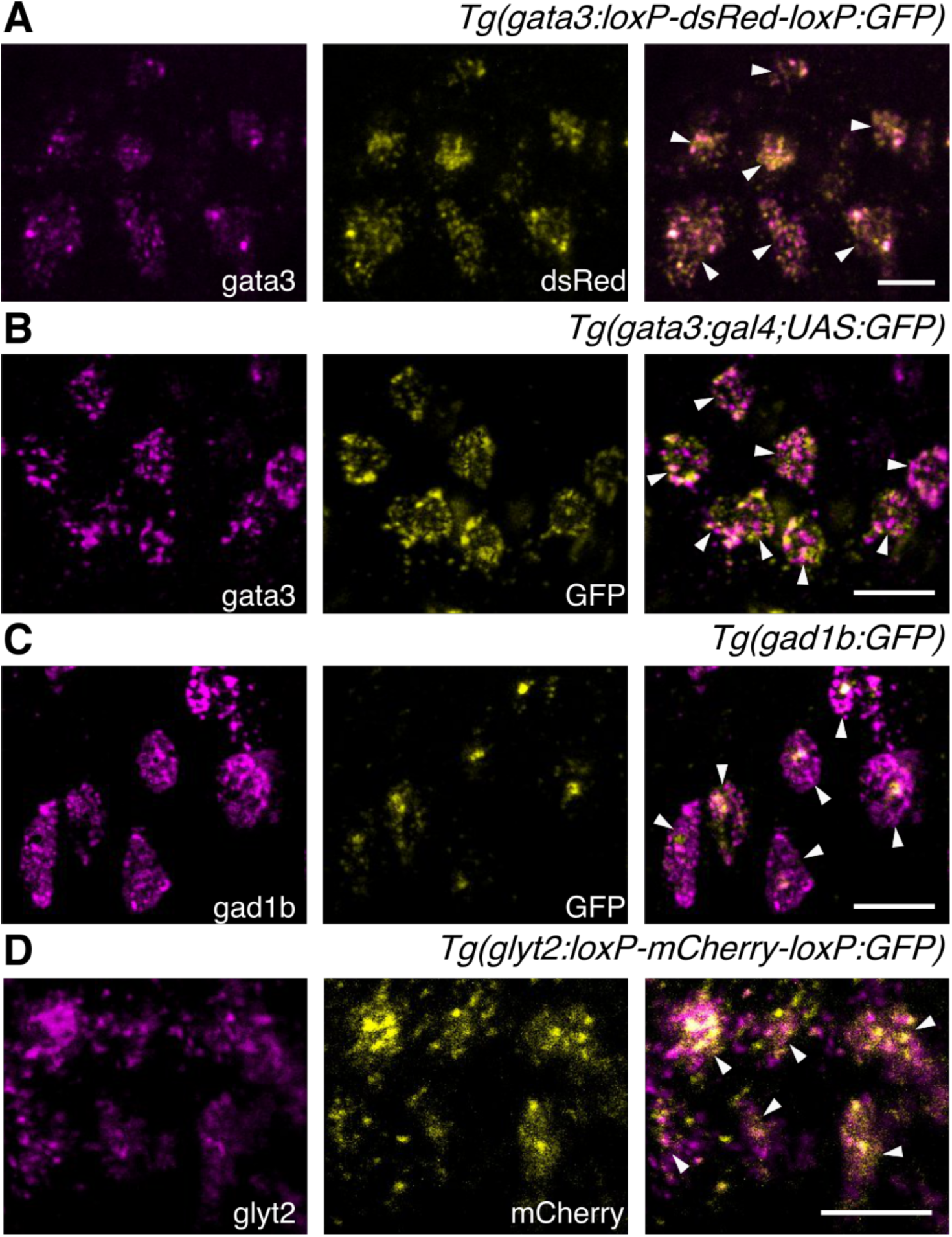
Two-color fluorescent in-situ hybridization validates transgenic line expression patterns. Confocal images (z-projection of ∼5 µm) showing fluorescent in-situ hybridization for endogenous RNA (magenta, left), transgenic fluorophore (yellow, middle) and overlaid two-color image (right). White arrowheads indicate colocalization. (A) *Tg(gata3:loxP-dsRed-loxP:GFP)*; (B) *Tg(gata3:gal4,UAS:GFP)*; (C) *Tg(gad1b:GFP)*; (D) *Tg(glyt2:loxP-mCherry-loxP:GFP)*. Scale bars = 10 μm.

**Table 1.** Summary of in-situ hybridization transgenic line validation, including completeness and accuracy.

|  | Completeness (%) | sd | Accuracy (%) | sd |
| --- | --- | --- | --- | --- |
| *Tg(gata3:loxP-DsRed-loxP:GFP)* | 96.57 | 5.34 | 95.75 | 6.45 |
| *Tg(gata3:gal4;UAS:GFP)* | 84.36 | 10.79 | 89.64 | 5.92 |
| *Tg(gad1b:GFP)* | 93.33 | 6.67 | 88.77 | 12.69 |
| *Tg(glyt2:loxP-mCherry-loxP:GFP)* | 86.51 | 7.26 | 92.21 | 2.70 |

### Neurotransmitter expression defines subpopulations of V2b neurons

Previous work has established that V2b neurons in embryonic zebrafish, as identified by Gata3 RNA expression, are exclusively inhibitory and predominantly GABAergic [24]. However, some spinal neurons are known to switch inhibitory neurotransmitters at early developmental stages [36, 37]. To resolve the neurotransmitter profile of V2b neurons in larvae, Gata3+ neurons were evaluated for coexpression with transgenic markers for *glyt2*, a glycine transport protein, and *gad1b*, a GABA synthesis enzyme. Nearly all larval V2b neurons expressed Glyt2 in 5 dpf larvae (Fig. 3A), in contrast to embryonic stages. Furthermore, Gad1b is expressed in approximately half of the V2b population (Fig. 3B).

**Figure 3.**
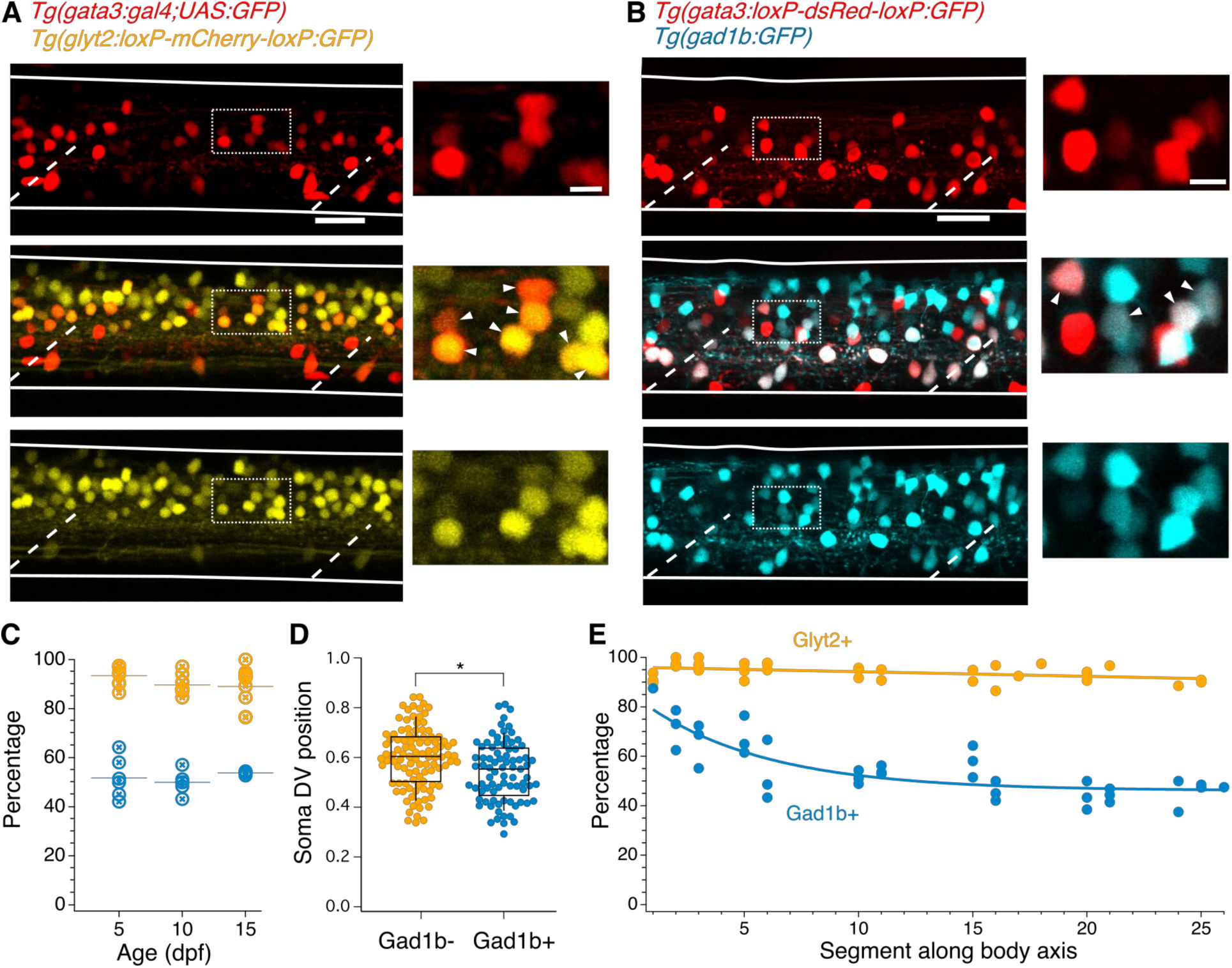
V2b neurons express the inhibitory neurotransmitter markers Glyt2 and Gad1b. (A) Lateral z-projection of a spinal cord hemisegment in a *Tg(gata3:gal,UAS:GFP;glyt2:loxP-mCherry-loxP:GFP)* (gata3, top; glyt2, bottom) double transgenic animal with composite image (middle). Dashed lines indicate muscle segments and solid lines indicate the spinal cord dorsal and ventral boundaries. Magnified inset, from dashed box, showing soma-level colocalization is shown to the right. Soma colocalization indicated with white arrowheads. Scale bar = 20 μm; inset 5 μm. (B) *Tg(gata3:loxP-DsRed-loxP:GFP;gad1b:GFP)* (gata3, top; gad1b, bottom) and dual-color composite image (middle). Magnified inset, from dashed box, is shown to the right. Soma colocalization indicated with white arrowheads. Scale bar = 20 μm; inset 5 μm. (C) Percentage of V2b neurons co-expressing GlyT2 or Gad1b is stable from ages 5-15 dpf, as measured in body segments 15-16. N = 6 animals at each time point. (D) V2b soma position for Gad+ and Gad-neurons differs slightly in the dorsoventral axis, *p < 0.01, Student’s t-test. (E) Percentage of V2b neurons co-expressing GlyT2 or Gad1b along the rostrocaudal body axis.

Inhibitory neurotransmitter switching is posited to occur at early developmental stages in zebrafish [36]. Therefore, we examined whether the variation of neurotransmitter expression in V2b neurons at 5 dpf represented a transient developmental stage or a stable pattern of expression. We assessed coexpression of the neurotransmitter markers in midbody V2b neurons at 5, 10, and 15 dpf, after which V2b neurons are not reliably labeled by transgenic lines (data not shown). Gad1b and Glyt2 expression in V2b neurons remains unchanged across these ages, with ∼52% of neurons expressing Gad1b and ∼91% expressing GlyT2 (Fig. 3C).

Are GABAergic and non-GABAergic neurons distributed similarly throughout the neuraxis? By plotting dorsal-ventral (DV) position relative to spinal boundaries, we found that on average, GABAergic V2b somata are located slightly ventral to non-GABAergic V2b somata, but that both populations span the same DV range (Fig. 3D). Therefore, soma position is not predictive of neurotransmitter expression. In the rostrocaudal axis, the percentage of GABAergic V2b cells is highest (∼80%) in rostral segments, then decreases to ∼50% by midbody and throughout the rest of the spinal cord. In contrast, Glyt2 robustly colabels with V2b cells throughout the entire spinal cord (Fig. 3E). These data indicate that the Gad1b+ and Gad1b-populations comprise distinct and persistent sunclasses. V2b neurons expressing both Glyt2 and Gad1b will be referred to as V2b-mixed, in reference to their mixed neurotransmitter expression, whereas V2b neurons that solely express Glyt2 will be referred to as V2b-gly.

### Axonal morphology varies by subpopulation identity

The classic axonal morphology of zebrafish VeLD neurons is ventral, with little change in DV position from the onset [24]. However, some V2b neuron fills exhibited axons with much more dorsal trajectories (e.g. Fig. 1D). To resolve whether these represent different subclasses, we investigated axonal morphology of identified V2b-mixed and V2b-gly neurons using single-cell dye electroporation in the double transgenic line *Tg(gata3:loxP-DsRed-loxP:GFP; gad1b:GFP)*, in which expression of GFP (Gad1b) differentiates between the mixed and glycinergic subclasses.

Although both V2b-gly and V2b-mixed neurons extend axons caudally and ipsilaterally, consistent with data in Fig. 1, the DV position of their axons was different. GABAergic V2b-mixed neurons projected axons ventrally along the spinal cord, with an average axon location found between 0.24 – 0.33 DV (example, Fig. 4A). In contrast, axons from V2b-gly neurons typically make an initial ventral dip but then turn more dorsally, ranging from 0.31 – 0.65 in the DV axis (example, Fig. 4B). Traces from all filled neurons are shown in Fig. 4C, and averaged trajectories in Fig. 4D. Somata were filled in segments ranging from 14 to 18; the traces are shown aligned at the soma for ease of visualization.

**Figure 4.**
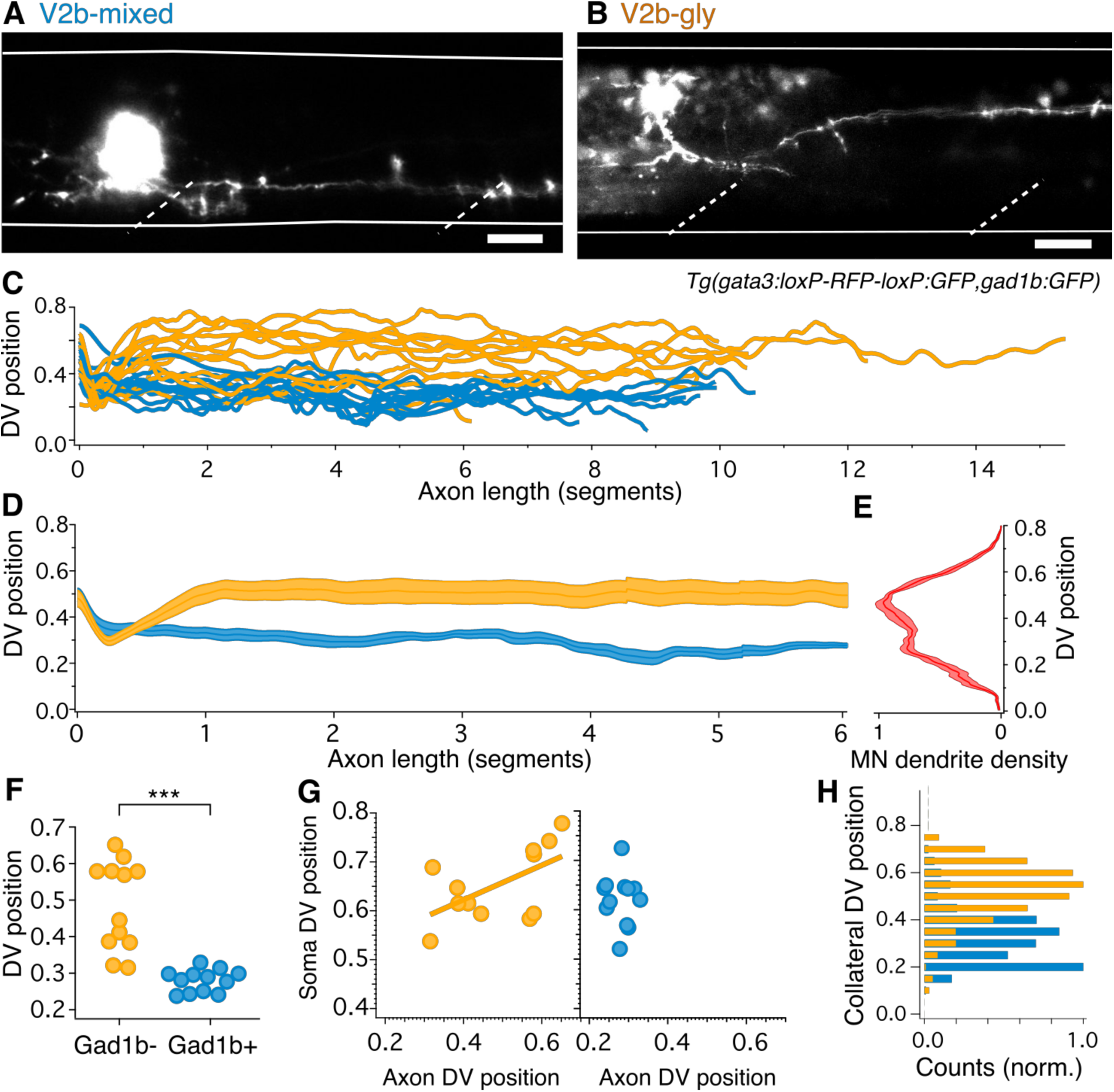
V2b-gly and V2b-mixed V2b neurons have distinct axon morphology and innervation territories. (A) Examples of a V2b-mixed (*Tg(gad1b:GFP)*+) and a (B) V2b-gly (*Tg(gad1b:GFP)*-) single-cell dye fill. Dashed lines indicate muscle segments and solid lines indicate the spinal cord dorsal and ventral boundaries. Scale bars = 20 μm. (C) Axon traces for V2b neurons, aligned at the segment of origin, relative to the spinal cord dorsoventral boundaries (V2b-mixed, cyan, n = 12; V2b-gly, orange, n = 12). All axons were exclusively descending. (D) Mean and SEM of V2b-gly and V2b-mixed axon trajectories. (E) Motor neuron dendrite fluorescence intensity, measured in *Tg(mnx:GFP)*, relative to the same dorsoventral landmarks. (F) Mean axon position for each traced axon. ***p < 0.0001, Student’s t-test. (G) Average axon position of V2b-mixed (cyan, left) and V2b-gly (orange, right) relative to soma position for each neuron. A correlation between soma position and axon position is observed for V2b-gly but not V2b-mixed neurons. V2b-gly: r^2^ = 0.33, p < 0.05, V2b-mixed: r^2^ = 0.0059, p = n.s. (H) Axon collaterals of V2b-gly neurons also innervate more dorsal spinal cord territory than V2b-mixed axons.

Other features of anatomy also varied between the two subtypes. The axon DV position of V2b-gly neurons is positively correlated to the soma DV position, i.e. a more dorsal soma projects a more dorsally positioned axon (Fig. 4G). However, this trend is not realized for V2b-mixed cells, which project axons ventrally to a narrow spinal cord region regardless of soma position. Putative en passant boutons were found in both cell populations. Most filled axons (22/24) extended vertical collaterals from the main axon. The number of collaterals per axon did not significantly vary between populations (V2b-mixed, median = 3, range 0-5; V2b-gly, median = 5, range 0-23, Mann-Whitney Wilcoxon test p = 0.056). However, collaterals of V2b-mixed and V2b-gly axons cover largely distinct DV regions of the spinal cord (Fig. 4H).

What is the significance of differential DV axon trajectories between V2b-gly and V2b-mixed subclasses? Previous work has shown that motor neurons active during fast movements are located more dorsally within the spinal cord, whereas those for slower movements are located more ventrally [21]. Therefore, we compared population averages of the V2b-gly and V2b-mixed axons (Fig. 4D) to a plot of motor neuron dendritic territory (Fig. 4E; see Methods) shown in Figure 4D. Notably, V2b axon position of the two classes overlaps with two peaks in the motor neuron density profile. Consequently, we next investigated the direct influence of V2b neurons on motor neurons.

### V2b subpopulations provide differential inputs to fast and slow circuits

Anatomical evidence indicates that V2b neurons make contact onto limb motor neurons where they are partially responsible for enforcing flexor/extensor alternation [18, 19]. However, to date there are no physiological recordings of synaptic connections from V2bs to other neurons in any species. We first validated that optogenetic stimulation in the *Tg(gata3:Gal4; UAS:CatCh)* line was sufficient to elicit action potentials in V2b neurons (Figs. 5A and 5B). We then targeted spinal motor neurons for *in vivo* recording in *Tg(gata3:Gal4, UAS:CatCh)* larvae at 4-6 dpf (Fig. 5C). Optogenetic activation of V2b neurons with a 20-50 ms pulse of light delivered 3-7 segments rostral to the recording site elicited robust IPSCs in motor neurons (Fig. 5D). Synaptic conductance amplitudes exhibited a median of 139 pS (25-75% range, 97-174 pS). Although the *Tg(gata3:Gal4)* line labels CSF-cNs in addition to V2bs (Fig. 1B), CSF-cNs exhibit short ascending axons that do not contact motor neurons other than the CaP [38]. To validate that V2b neurons are providing these inhibitory inputs, we used a digital micromirror device to deliver targeted squares of light stimuli (∼20 μm × 20 μm) to dorsal spinal cord areas containing V2b but not CSF-cN somata. These localized stimuli still elicited reliable IPSCs in both primary and secondary motor neurons (Figs. S1B and S1C). As a second control, recordings were also made in a subset of animals with strong CatCh expression in CSF-cNs and negligible expression in V2b neurons (see Methods and Fig. S1D). In these recordings, even full-field light stimulation evoked only minimal IPSCs in motor neurons (Fig. S1E). Together these results indicate that the optogenetically elicited inhibitory inputs arise from monosynaptic V2b to motor neuron connections.

**Figure 5.**
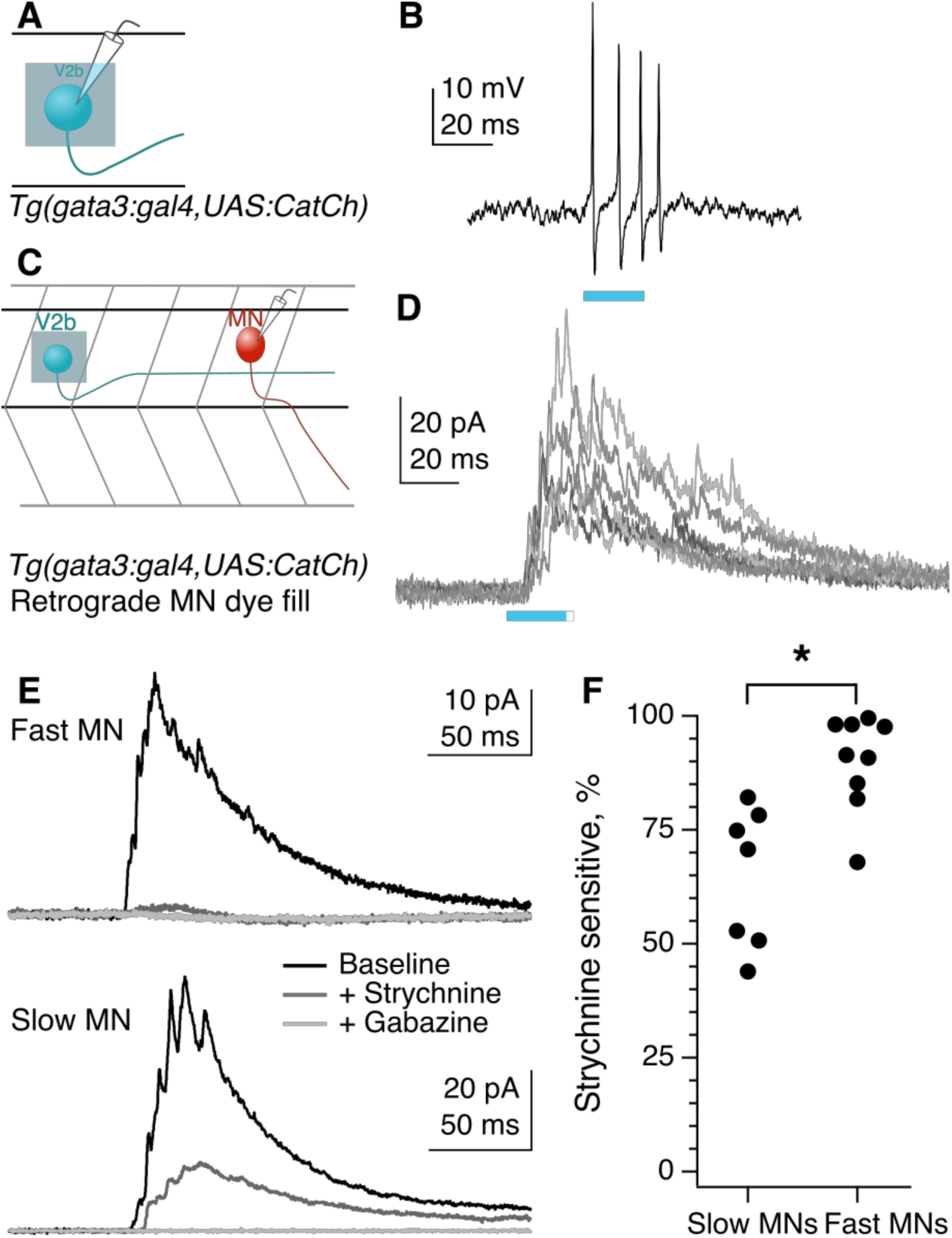
Fast motor neurons receive predominantly glycinergic V2b inputs, whereas V2b synaptic inputs to slow motor neurons are mediated by both GABA and glycine receptors. (A) Schematic of recording to validate CatCh expression in V2b neurons. (B) Cell-attached recording from a V2b neuron expressing CatCh during a 20 ms illumination epoch. Note that evoked action potentials outlast the duration of illumination, presumably due to membrane depolarization and/or Ca influx. (C) Schematic illustrating whole-cell recordings from motor neurons paired with optogenetic stimulation of V2b neurons. (D) Six overlaid sweeps showing ISPCs barrages recorded in a motor neuron in response to optogenetic activation of V2b neurons. Blue bar represents the light stimulus. All recordings were carried out in the presence of NBQX. (E) Average IPSC responses to light stimulation in fast (top) and slow (bottom) motor neurons, as identified by soma location and input resistance. Response during baseline (black), after application of strychnine (dark grey), and after additional application of gabazine (light grey). In all cases, the IPSC was entirely abolished by the combination of strychnine and gabazine. (F) Percentage peak current reduction by strychnine in fast and slow motor neurons. *p < 0.01.

The striking difference in dorsal-ventral targeting of V2b-gly and V2b-mixed axonal trajectories (Fig. 4D) suggests a potential relationship with the well-described dorsal-ventral distribution of motor neurons based on size and speed at recruitment. Large motor neurons with low input resistance are located dorsally within the motor pool and are recruited for the fastest speeds of swimming, whereas more ventrally located motor neuron somata exhibit higher input resistance and are recruited during slower movements [21, 22]. Accordingly, we tested whether the glycinergic and GABAergic components of the IPSC differed between fast and slow motor neurons. Bath application of strychnine to block glycine receptors abolished a median of 91% of the V2b-evoked IPSC in fast motor neurons, but only 71% of the V2b-evoked IPSC in slow motor neurons (Fig 6F, C; p = 0.003, Wilcoxon Rank test). The GABA_A_ receptor antagonist gabazine (SR-95531) eliminated the remaining IPSC in all cases. Therefore V2b-mediated inhibition onto fast motor neurons is carried out predominantly by glycinergic synapses, whereas V2b inhibition onto slow motor neurons is carried by mixed glycinergic/GABAergic transmission. These results are consistent with the idea that V2b-gly preferentially inhibit more dorsally located fast motor neurons, whereas V2b-mixed inhibit the more ventrally located slow motor neurons.

**Figure 6.**
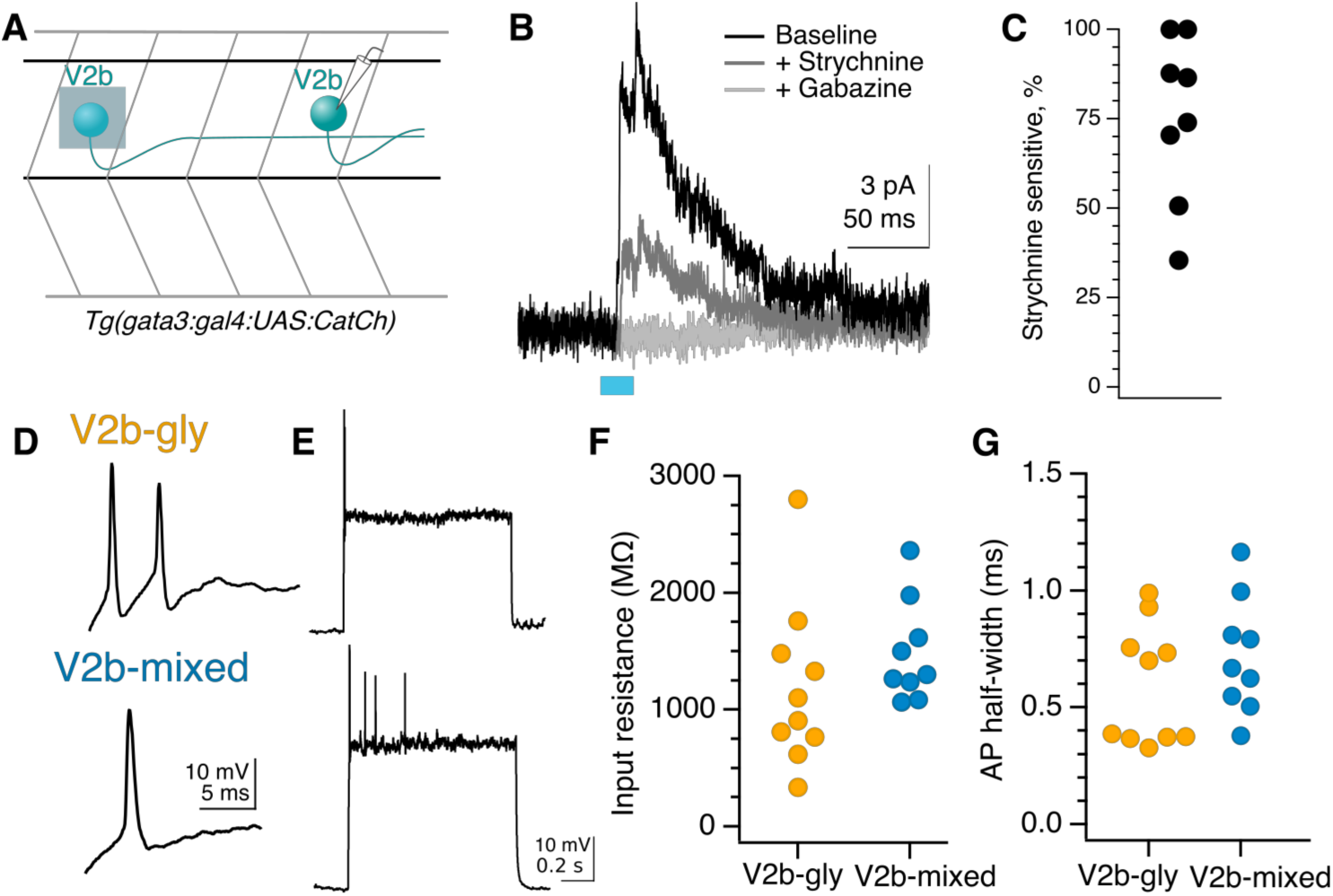
Rostral V2b neurons inhibit more caudal V2b neurons, providing circuit disinhibition; V2b-gly and V2b mixed populations are physiologically indistinguishable. (A) Example action potential magnified from (B) responses to step depolarizations in both classes of V2b neurons. Most recorded neurons in both groups could not sustain action potentials across a step. (C) Input resistance measured via hyperpolarizing test pulse. N = 10 Gad− (orange), 9 Gad+ (cyan). (D) Action potential peak half-widths are not significantly different between the two groups. (E) Experimental schematic for V2b-to-V2b connectivity recordings. (F) Evoked IPSCs recorded in an example V2b neuron in response to optogenetic stimulation of more rostral V2b neurons, black, and the response after the successive addition of glycine and GABA_A_ receptor antagonists, dark grey and light grey traces respectively. The blue bar represents the duration of optogenetic stimulation. (G) Percentage peak current sensitivity to strychnine.

Some spinal premotor neurons form synaptic connections within their own populations, suggestive of speed- or state-related “gears” [39]. Optogenetic activation of rostrally-located V2b neurons evoked IPSCs in 11/21 (52%) mid-body V2b neurons (Figs. 6A and 6B). The median conductance of individual IPSCs was 158 pS (25-75% range, 84-158 pS). Application of strychnine blocked a median of 80% of the V2b-evoked IPSC, while the remainder was abolished by gabazine (Figs. 6B and 6C). Thus, some V2b neurons inhibit other members of the V2b pool forming a disinhibitory pathway.

### V2b cell physiology does not distinguish between subtypes

Intrinsic physiological characteristics, including input resistance and spiking properties, can be used to subdivide some spinal interneuron populations into distinct subpopulations [2, 6, 40]. We examined whether the V2b-gly and V2b-mixed subgroups exhibited differences in intrinsic physiology by targeting whole-cell recordings to these neurons. V2b neurons were silent at rest, in contrast to CSF-cNs which exhibited spontaneous spiking (data not shown). Spikes were elicited by depolarizing current steps, (Figs. 6D and 6E) which usually led to one or a few spikes, with only 3/10 V2b-gly and 3/9 V2b-mixed neurons able to sustain spiking across the step. There was no difference in input resistance (Fig. 6F) or spike shape (Fig. 6G) between the V2b-gly and V2b-mixed neurons. Therefore, the two V2b subpopulations are indistinguishable at the level of intrinsic physiology despite their differences in axon trajectory.

### Optogenetic V2b suppression increases tail beat frequency

What are the functional consequences of V2b inhibition onto motor neurons (Fig. 5)? To better understand this role, we carried out high-speed behavioral recordings during optogenetic inactivation of V2b neurons with a light-gated Cl^−^ channel, ZipACR [41]. To eliminate contributions from Gata3 expressing neurons in the brain, we used a spinally transected preparation. Tail movements were induced pharmacologically with application of N-methyl-d-aspartate (NMDA, 200 μM) [42]. NMDA induces tail movements with episodic, left-right alternations that mimic the natural beat-and-glide swims of 5 dpf larvae [42–44].

Spinal CSF-cNs are labeled in BAC-generated Gata3 transgenic lines (Fig. 1A). However, a CRISPR-generated *Tg(gata3:zipACR-YFP)* knock-in shows robust expression of the fluorescent ZipACR protein in V2b neurons but only sparse, dim expression in CSF-cNs (Fig 7A). Within CSF-cNs expression was observed exclusively in the apical extension into the central canal but not the soma (Figs. 7A and S2). We first validated the efficacy of the ZipACR construct in suppressing V2b firing under high-intensity light (Fig. S3A, n = 4). Under lower-intensity light conditions, identical to those of the behavioral recordings, action potentials were completely suppressed in 4 out of 6 V2b cells and partially suppressed in 1 additional cell (Figs. 7B and C), in *Tg(gata3:zipACR-YFP; gata3:loxP-DsRed-loxP:GFP)* animals. In contrast, identical stimulation partially suppressed spiking in only 1 of 5 CSF-cNs (Figs. 7C and S3C). Therefore, we used these light stimulation parameters, under which V2b neurons are mostly if not entirely suppressed whereas CSF-cNs are not substantially affected, to carry out behavioral experiments assessing the effects of suppressing V2b neurons on locomotion.

**Figure 7.**
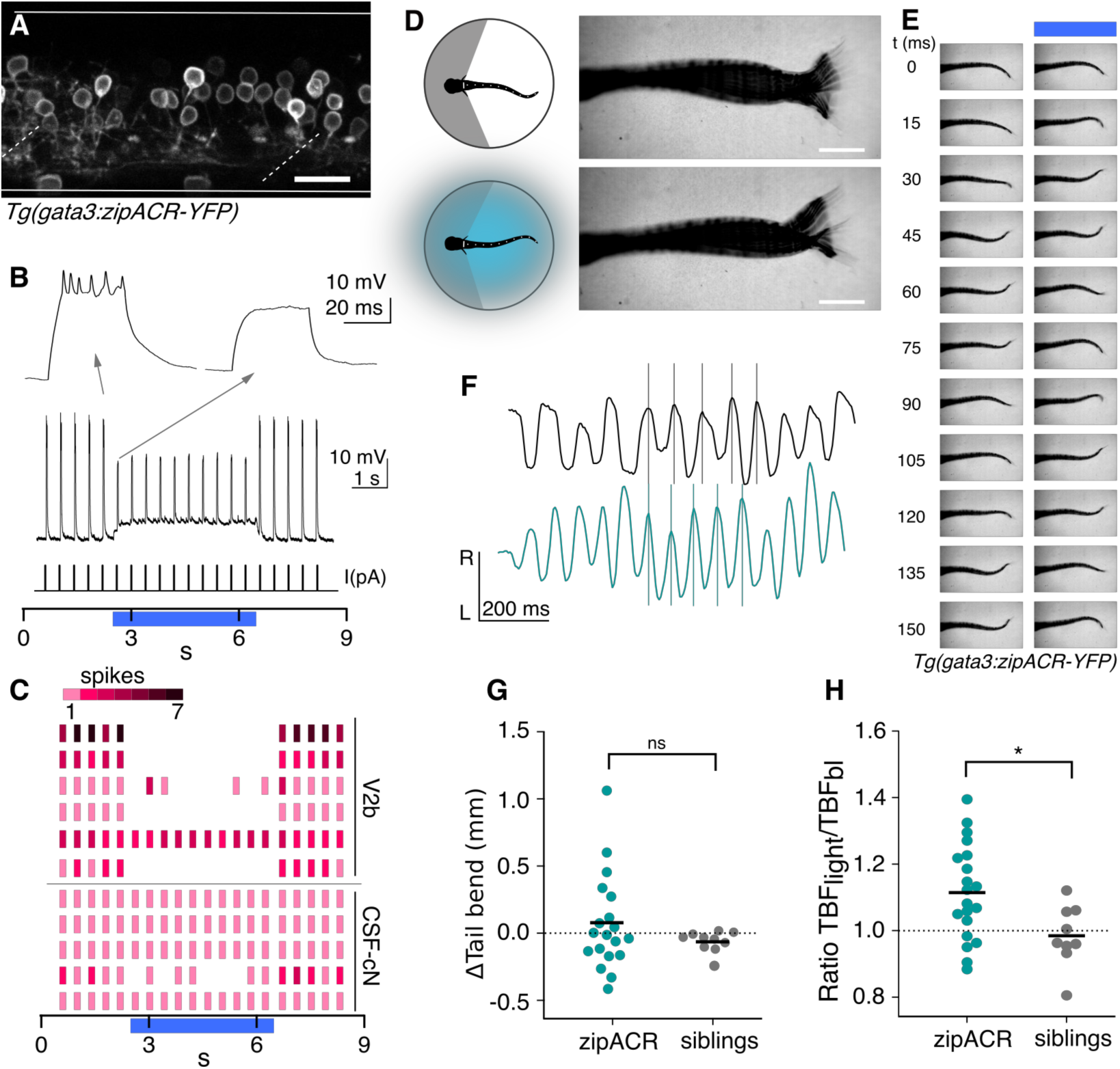
Optogenetic suppression of V2b activity leads to increased locomotor speeds. (A) Z-projection of *Tg(gata3:zipACR-YFP)* over one full segment of spinal cord showing expression in V2b but not CSF-cN somata. CSF-cN apical extentions show minimal YFP expression. See also Fig. S2. Scale bar= 20 μm. (B) A whole cell recording during repeated current steps (20 ms duration) <u>is</u> shown for an example V2b neuron in a *Tg(gata3:zipACR-YFP)* animal. Blue bar indicates period of optical stimulation. An expanded view of current steps before and during optical stimulation are shown above with arrows. Recordings indicate that current steps normally elicit bursts of action potentials, but coincident optogenetic suppression prevents spiking, yielding only subthreshold depolarizations. (C) Raster plot of action potentials for 6 V2b cells and 5 CSF-cN cells summarizes optogenetic suppression across cell types. Color value represents number of spikes elicited during each current step. 5/6 V2b neurons were mostly or entirely suppressed, whereas only 1/5 CSF-cN were affected. (D) Schematic of behavioral recording depicting the NMDA-induced tail movements of spinalized head-embedded animals without and with optogenetic stimulation. Image overlay of 100 ms of tail movements without and with light stimulation in a *Tg(gata3:zipACR-YFP)* animal show striking similarities in tail displacement during swim. Scale bar = 0.5 mm. (E) Time point images of tail movements in the same animal at baseline (left) and with optogenetic stimulus (right) demonstrating the difference in timing of tail movements during V2b suppression. (F) Tracked left-right tail position during recordings with (teal) and without (black) optical stimulation for the same *Tg(gata3:zipACR-YFP)* animal. Lines for each recording are aligned to consecutive peaks in the baseline trace to illustrate the phase advance and increased tail beat frequency during optogenetic stimulation (teal). (G) Average change in tail bend between stimulation and control recordings during swim movements for each animal, ns p = 0.14. (H) Ratio of average TBF during stimulation to baseline TBF for each animal, cohort averages shown with black dash. N = 20 *Tg(gata3:zipACR-YFP)* and N = 9 siblings. *p < 0.01.

Animals were head-embedded with a free tail and high-speed (200 Hz) recordings were acquired to capture fictive locomotion. Swim dynamics were recorded and evaluated both with and without optogenetic stimulation (Figs. 7D and 7E). Kinematic analysis of the high-speed video was performed with code adopted from [45]. The total tail displacement and quantity of tail movements did not significantly differ during optogenetic stimulation (Figs. 7D and 7G).

Strikingly, tail beat frequency (TBF), a metric of locomotor speed, increased in *Tg(gata3:zipACR-YFP)*+ animals during light stimulation but not in their ZipACR negative clutchmates (Figs. 7E, 7F, and 7H, paired t-test p = 0.0025). The average TBF change was 1.4 Hz and was robustly observed in animals from three clutches. In contrast, elimination or genetic silencing of CSF-cN causes the opposite effect, a decrease in TBF [46]. Therefore, any modest changes in CSF-cN activity in our experiments (Fig. 7C) would if anything serve to counteract the effects of V2b silencing. We conclude that suppression of V2b neurons increases TBF, and consequently the normal role of V2b-mediated inhibition is to serve as a brake on locomotor frequency.

## Discussion

In this study we demonstrated that V2b neurons exert direct control over axial musculature in the larval zebrafish. The V2b population comprises two stable subclasses, defined by neurotransmitter identity: one subclass is exclusively glycinergic and the other mixed glycinergic/GABAergic. These distinct V2b-gly and V2b-mixed subclasses preferentially inhibit fast and slow motor neurons, respectively, analogous to the speed-dependent connectivity found in a diverse range of zebrafish spinal interneurons [23, 31, 47, 48]. Moreover, we found that the suppression of V2b activity led to an increase in locomotor speeds. Together, these results indicate that inhibition from V2b neurons is not restricted to enforcing agonist-antagonist muscle coordination but also influences locomotor speed through in-phase modulation of axial motor neurons.

### V2b conservation across species

We demonstrated that V2b neurons are inhibitory and extend axons ipsilaterally and caudally in zebrafish (Figs. 1, 3) similar to V2b neurons in mice [18, 25]. Gata3+ V2b neurons are widely present in vertebrates but have also been identified in the nerve cord of a marine annelid indicating an ancestral persistence in motor circuitry [32, 49]. V2b neurotransmitter profiles appear to vary throughout development and across species. Gata3-expressing cells in the embryonic 24 hours post fertilization zebrafish are predominantly GABAergic with a smaller subset expressing or co-expressing glycine [24], an inversion of our finding that V2b cells in 5-15 dpf zebrafish are all glycinergic with approximately half co-expressing GABA. Early in development, murine V2b neurons broadly co-express GABA and glycine [19]. By P0 in mouse, however, nearly all V2bs are glycinergic and ∼25% are GABAergic [25], which is broadly similar to our results. In zebrafish, these two subclasses persist out to 15 dpf which implies that they are distinct identities.

Consistent with the idea that V2b-gly and V2b-mixed are distinct identities, the two subclasses exhibit different axon trajectories in the DV axis, perhaps indicating responsiveness to different axon guidance cues. In mouse, V2b subpopulations have not been directly shown. However, the differential expression of the transcription factors Gata2/Gata3/BhlhB5 in non-overlapping neural subsets may imply their presence [50]. More broadly, our finding of subclasses in the V2b population is parallel to previously identified genetically and anatomically distinct subclasses within the V0, V1 and V2a populations in the mouse and zebrafish [2, 4-6, 51].

### Do V2b-gly and V2b-mixed populations match existing zebrafish neural classes?

Historically, zebrafish spinal neurons have been classified by anatomy [52]. V2b (Gata3+) neurons are thought to correlate to ventral longitudinal (VeLD) neurons, an anatomically defined cell class with a characteristic longitudinal ventral-positioned axon [24, 53]. How do the subpopulations we have described here relate to the VeLD population? Based on the ventral axon morphology and GABA co-expression, the V2b-mixed subtype represents a matured version of the embryonic VeLD neurons. In contrast, V2b-gly neurons are distinct in morphology and neurotransmitter profile from VeLDs, indicating either that they have not been characterized in embryonic stages or that they develop at a later time.

In mice, the V2 progenitor domain gives rise to a third class of neurons called V2c, which express Sox1 and only transiently Gata3 in very early development (prior to E12.5) before later downregulation [54]. It is unclear whether zebrafish have a homologous V2c population although Sox1a/b is present in the 24 hpf spinal cord and notably also colabels with Gata3 [55]. A possible V2c homolog, referred to as V2s, has been identified as a Sox1a/b+ glycinergic cell type deriving from the V2 domain with long, ipsilateral, caudally projecting axons [56] similar to the V2b-gly neurons described here. However, Gata3 expression was not investigated in V2s neurons, leaving it unclear whether V2s neurons are in fact V2b-gly [56]. Given the persistent, distinguishing expression of Gata3 in both V2b-gly and V2b-mixed subtypes, our data are consistent with the designation of two subclasses within V2b, not a V2c homolog or additional V2s class. Further detailed investigation of Sox1a/b gene expression in these neurons will be required to clearly separate these classes.

### Speed specific inputs to motor circuits

Locomotion at faster versus slower speeds engages different sets of spinal interneurons, both within a genetically defined population [23, 29, 31, 47] and across populations [21, 57]. Given the observation that slow motor neurons likely receive more input from V2b-mixed neurons whereas fast motor neurons receive largely V2b-gly input, it would be of interest to explore whether the V2b subpopulations are recruited at different speeds of locomotion. Furthermore, the intra-V2b connectivity suggests a possible “gear shift” within the V2bs, with the V2b-gly and V2b-mixed populations potentially inhibiting each other to enforce a given speed of swim. One caveat in interpretation of these results is that despite their different somatic positions, fast and slow motor neurons have overlapping dendritic fields [48]. Therefore, it is possible that V2b-mixed neurons make synapses onto all motor neurons, and differential receptor expression is responsible for their IPSC pharmacology (Fig. 5); meanwhile, V2b-gly might be responsible for other functions, such as suppression of dorsal horn sensory interneurons [58]. Paired recordings or higher resolution anatomical experiments will be required to distinguish these possibilities.

### V2b suppression increases tail movement speed

One role of ipsilateral inhibition is to mediate flexor-extensor alternations via Ia reciprocal inhibition from V2b and V1 populations [18, 19]. Ipsilaterally descending propriospinal neurons may also stabilize left-right alternation, although specific ablation of inhibitory ipsilaterally descending neurons has not been tested [8, 59]. Our work establishes that selective suppression of V2b activity increases tail beat frequency (Fig. 7). In contrast, genetic ablation of V1 neurons in mouse dramatically reduced fictive step speed [60]. Similarly, pertubation of in-phase inhibition led to slower crawling speeds in larval Drosophila and reduced rhythmic motor drive in Xenopus [13, 61]. From this we surmise that ipsilateral inhibition from V1 and V2b shape distinct features of locomotion. V1 neurons may act to terminate motor neuron burst cycles while V2b neurons may limit overall speed, much like a brake.

The behavioral outcome of V2b inactivation is modest at slow speeds. However, V2a influence over motor circuits strengthens for increasing speeds [21, 47]. Thus, it is possible that the effect of silencing V2b neurons will be larger at faster locomotor speeds, particularly given that in-phase inhibition increases for faster movements [10, 11, 62]. It is unclear from current experiments whether the V2b-associated speed increase is due to the loss of on-cycle V2b inhibition onto motor neurons or through inputs to the premotor circuitry, for example through V2b-V2b interconnectivity (Fig. 6). More selective optogenetic manipulations will be useful to separate out these effects.

### Overall role of V2b in motor circuits

What is the functional role of V2b-mediated ipsilateral inhibition onto motor circuits? Three broad categories of V2b function occur. First, as discussed above, enforcing flexor-extensor and potentially forelimb-hindlimb alternation in limbed animals. Second, as supported in this work, V2b neurons may serve to titrate motor neuron spiking differentially across varying speeds of movement. Measuring inhibitory conductances in motor neurons *in vivo* has revealed, surprisingly, that inhibition in-phase with excitation actually increases for increasingly strong movements [10, 11, 13, 62], rather than diminishing to allow more powerful contractions. In this context, shunting ipsilateral inhibition might serve to enforce tight temporal control over spike timing via shortening membrane time constants. Alternatively or in addition, ipsilateral inhibition might function as a form of gain control to prevent saturation, analogous to somatic inhibition in hippocampus and cortex [11, 63, 64].

Thirdly, ipsilateral inhibition may act to isolate movements in certain behaviors that engage dedicated premotor circuitry, e.g. through the selective inhibition of interneurons during scratching versus swimming in turtle [65, 66]. Some V2b neurons, by virtue of their direct connections with ipsilateral motor circuits, could form part of the locomotor “switch” from one behavior to another. A thorough investigation of V2b-gly and V2b-mixed recruitment during natural behaviors, such as speed transitions, turning, or balance, will allow us to better understand the similar or distinct ways that V2b subclasses influence locomotion.

## Methods

### Animal care

Adult zebrafish (Danio rerio) were maintained at 28.5°C with a 14:10 light:dark cycle in the Washington University Zebrafish Facility following standard care procedures. Larval zebrafish, 4-7 days post fertilization (dpf), were kept in petri dishes in system water or housed with system water flow. Animals older than 7 dpf were fed rotifers daily. All procedures described in this work adhere to NIH guidelines and received approval by the Washington University Institutional Animal Care and Use Committee.

### Line generation

The *Tg(gata3:gal4)* and *Tg(gata3:LoxP-dsRed-LoxP:GFP)* lines were generated via the bacterial artificial chromosome (BAC) transgenic technique [67], using BAC zK257H17. The Gal4 and LRL-GFP constructs are described in Kimura et al. [28] and Satou et al. [68], respectively. The *Tg(glyt2:LoxP-mCherry-LoxP:GFP)* line was generated with CRISPR/Cas9 genome targeting methods utilizing the short guide RNA, donor plasmid, and methods described in Kimura et al. [69]. *Tg(gata3:zipACR-YFP)* animals were generated with CRISPR/Cas9 techniques using a gata3 short guide, TAG GTG CGA GCA TTG AGC TGA C. The donor Mbait-hs-zipACR-YFP plasmid was made by subcloning ZipACR [41], obtained from Addgene, into a Mbait-hs-GFP plasmid with Gibson Assembly cloning methods. A UAS:Catch [70] construct containing tol2 transposons was microinjected along with tol2-transposase RNA into one-cell *Tg(gata3:gal4)* embryos to generate the *Tg(gata3:gal4;UAS:Catch)* line.

### Single-cell photoconversion

Fluorescent protein photoconversion was performed on anesthetized and embedded 5 dpf *Tg(gata3:gal4; UAS:kaede)* animals using an Olympus FV1200 microscope. Single-plane confocal images were continuously acquired to monitor conversion progress while 500 ms bursts of 405 nm light (100% intensity) were applied to an ROI ∼1/10^th^ the size of the targeted soma to elicit photoconversion. Animals were removed from agarose and allowed to recover in system water for 1-3 hours. After recovery, fish were anesthetized, embedded, and imaged as above. Tiled image stacks were acquired over an area ranging from the most rostral processes to the most caudal with a minimum of 10% area overlap between adjacent fields of view to aid the image stitching process.

### Single-cell dye electroporation

*Tg(gata3:LoxP-dsRed-LoxP:GFP; gad1b:GFP)* animals (5-6 dpf) were anesthetized in 0.02% MS-222 and three electroetched tungsten pins were placed through the notochord securing the animal to a Sylgard-lined 10 mm well dish. Forceps and an electroetched dissecting tool were used to remove skin and one segment of muscle fiber to expose the spinal cord. A pipette electrode filled with 10% Alexa Fluor 647 anionic dextran 10,000 MW (Invitrogen) in internal recording solution, was positioned to contact the soma of the transgenic-labeled target neuron. Dye was electroporated into the cell via one or more 500 ms, 100 Hz pulse trains (1 ms pulse width) at 2-5 V (A-M systems Isolated Pulse Stimulator Model 2100). Confocal imaging was performed as described above, after > 20 min for dye filling.

### Fluorescent hybridization chain reaction (HCR)

Animals were fixed at 5 dpf in 4% paraformaldehyde and *in situ* hybridization was performed according to the HCR v3.0 protocol [71] with noted modifications. Preparation, dehydration and rehydration steps 1 through 14 were replaced with steps 2.1.1 through 2.2.8 with a Heat Induced Antigen Retrieval (HIAR) option in place of Proteinase K treatment [72, 73]. *In situ* probes were designed and distributed by Molecular Technologies (Beckman Institute, Caltech) to target gata3, gad1b, glyt2 (slc6a5), DsRed, mCherry, and GFP. Samples were kept in 4x saline-sodium citrate solution at 4°C prior to imaging. Samples were mounted in Vectashield (Vector Laboratories) or low-melting point agarose (Camplex SeaPlaque Agarose, 1.2% in system water) and positioned laterally on a microscope slides with #1.5 coverslip glass.

### Confocal imaging

5-7 dpf larvae were anesthetized in 0.02% MS-222 and embedded in low-melting point agarose in a 10 mm FluoroDish (WPI). Images were acquired on an Olympus FV1200 Confocal microscope equipped with high sensitivity GaAsP detectors (filter cubes FV12-MHBY and FV12-MHYR), and a XLUMPLFLN-W 20x/0.95 NA water immersion objective. A transmitted light image was obtained along with laser scanning fluorescent images. Sequential scanning was used for multi-wavelength images. Z-steps in 3D image stacks range from 0.8-1.4 microns. Fluorescent *in situ* hybridization samples were imaged with an UPLSAPO-S 30x/1.05 NA and silicone immersion media. Spectral images were collected for *Tg(gata3:zipACR-YFP; gata3:loxP-DsRed-loxP:GFP)* animals to distinguish between expression patterns of overlapping fluorophores. Samples were excited with a 515 nm laser. Emission was collected with a PMT detector from 10 nm wide spectral windows across the emission range 525-625 nm for each z-plane. Spectral deconvolution was performed with Olympus Fluoview software.

### Electrophysiology

Whole-cell patch-clamp recordings were targeted to V2bs or motor neurons in *Tg(gata3:gal4; UAS:CatCh), Tg(gata3:zipACR-YFP)*, doubly-transgenic *Tg(gata3:LoxP-dsRed-LoxP:GFP; gad1b:GFP) or Tg(gata3:zipACR-YFP; gata3:loxP-DsRed-loxP:GFP)* larvae at 4-6 dpf. Larvae were immobilized with 0.1% α-bungarotoxin and fixed to a Sylgard lined petri dish with custom-sharpened tungsten pins. One muscle segment overlaying the spinal cord was removed at the mid-body level (segments 9-13). The larva was then transferred to a microscope (Scientifica SliceScope Pro or Nikon Eclipse) equipped with infrared differential interference contrast optics, epifluorescence, and immersion objectives (Olympus: 40X, 0.8 NA; Nikon: 60X, 1.0 NA). The bath solution consisted of (in mM): 134 NaCl, 2.9 KCl, 1.2 MgCl_2_, 10 HEPES, 10 glucose, 2.1 CaCl_2_. Osmolarity was adjusted to ∼295 mOsm and pH to 7.5.

Patch pipettes (5-15 MΩ) were filled with internal solution for current clamp composed of (in mM): 125 K gluconate, 2 MgCl_2_, 4 KCl, 10 HEPES, 10 EGTA, 4 Na_2_ATP, 0.05-0.1 Alexa Fluor 647 hydrazide; for voltage clamp, 122 cesium methanesulfonate, 1 tetraethylammonium-Cl, 3 MgCl_2_, 1 QX-314 Cl, 10 HEPES, 10 EGTA, 4 Na_2_ATP and 0.05 – 0.1 Alexa Fluor 568 or 647 hydrazide. Osmolarity was adjusted to ∼285 mOsm and KOH or CsOH, respectively was used to bring the pH to 7.5. Recordings were made in whole-cell configuration using a Multiclamp 700B, filtered at 10 kHz (current clamp) or 2 kHz (voltage clamp) and digitized at 20 kHz with a Digidata 1440 or 1550 (Axon Instruments) and acquired with WinWCP (J. Dempster, University of Strathclyde).

The *Tg(gata3:gal4; UAS:CatCh)* line labels both V2b and Kolmer-Agduhr / cerebrospinal fluid-contacting neurons (CSF-cNs). To ensure that evoked IPSCs derived from presynaptic V2bs rather than CSF-cNs, epifluorescent illumination was targeted 3-7 segments rostral to the recorded segment. A Polygon400 Digital Micromirror Device (Mightex) was used to provide patterned illumination in indicated recordings. Previous studies found that CSF-cNs have short ascending axons and do not contact any motor neurons besides the caudal primary (CaP) [38]. CatCh expression in the *Tg(gata3:Gal4; UAS:CatCh)* line is variegated, with some animals showing strong CatCh expression throughout both CSF-cNs and V2bs, and others showing good expression in CSF-cNs and minimal expression in V2b cells. Additional control experiments were performed in animals with minimal V2b label to demonstrate the absence of contribution of CSF-cN synapses in these experiments.

Motor neurons were identified by axon fill that extended into the musculature and/or by retrograde dye labeling from the muscle. For retrograde labeling, 4 dpf larvae were anesthetized (0.02% MS-222) and laid flat on an agarose plate. A Narishige micromanipulator in conjunction with a microinjection pump (WPI, Pneumatic Picopump) was used to deliver small volumes of dye (Alexa Fluor 568 dextran, 3000 MW) via glass pipette into the muscle. Fish recovered in regular system water and were subsequently used for recordings at 5-6 dpf.

Data were imported into Igor Pro using NeuroMatic [74]. Spike threshold was defined as 10 V/s, and custom code was written to determine spike width and afterhyperpolarization of the initial spike elicited by pulse steps. Input resistance was calculated by an average of small hyperpolarizing pulses. To isolate IPSCs, 10 µm NBQX was present in the bath and neurons were voltage clamped at the EPSC reversal potential.

Motor neurons at the dorsal extent of the distribution (> 50% of distance from bottom of spinal cord to top) exhibited lower input resistances (mean ± SD: 287±75 MΩ) and were considered “fast” and the remainder, which exhibited higher input resistances (885±367 MΩ) considered “slow” [21]. These groups mostly correspond to primary and secondary motor neurons, but some dorsally located bifurcating secondaries may be included in the fast group [22].

Optogenetic validation of ZipACR in V2b and CSF-cN was performed on *Tg(gata3:zipACR-YFP)* and *Tg(gata3:zipACR-YFP; gata3:loxP-DsRed-loxP:GFP)* animals. Light stimulation was provided with high intensity illumination, 5-10% intensity with a 40X (0.8 NA) water-immersion objective, and low intensity illumination which is identical to the conditions of behavioral recordings, 100% intensity with a 4X (0.1 NA) air objective.

### Image analysis

Image analysis was performed with ImageJ (FIJI) [75]. Igor Pro 6 was utilized for data analysis and statistics unless otherwise noted. V2b cell counts and neurotransmitter coexpression was quantified manually by two researchers (R.C. and M.J.); no significant differences in quantification were detected. Gata3+ V2b cells were identified and marked (ImageJ Cell Counter) relative to spinal cord and segment boundaries, giving total V2b/segment quantities. Subsequently, each cell was evaluated for expression of fluorescent proteins marking gad1b or glyt2.

Transgenic line validation was performed with in situ hybridization and quantified by two researchers (M.B. and R.C.) with no significant discrepancy in results. A ∼3-5 μm z-stack projection was made in a cell-dense area of spinal cord spanning two to three segments for each animal. ROIs of neurons were drawn in one channel before checking whether there was colocalization in the other channel. Samples were quantified twice: once for completeness (percentage of endogenous RNA positive neurons also expressing the transgene) and once for accuracy (percentage of transgene labeled neurons positive for endogenous RNA). 3-7 animals were evaluated in each line.

For axon tracing, stitched projection images were made with the Pairwise stitching [76] ImageJ plugin. The overlap of the fused image was smoothed with linear blending and was registered based on the fill channel or the average of all channels. Photoconversion cell fill images underwent an extra processing step in which the bleached green channel was subtracted from the photoconverted red channel. The Simple Neurite Tracer plugin [77] was used to trace the axon projection and branching relative to marked spinal cord boundaries. Axon lengths are reported as the number of segments transversed.

Motor neuron dendrites were quantified from confocal z-stack images of *Tg(mnx:GFP)* 5 dpf animals. Images were cropped to a single hemisegment. The Weka Trainable Segmentation plugin [78] was used to segment the motor neuron image into three classifiers; soma, axons exiting the spinal cord, and dendrites. Classification was based on Hessian training features. Training was performed iteratively for each image. The binary segmented images were applied to mask all non-dendrite fluorescence (n = 4 hemisegments/animal; n = 3 animals.) Fluorescence was maximum intensity projected in the z-dimension, collapsed along the horizontal plane and normalized to give an estimate of motor neuron dendrite density in the dorsoventral plane of the spinal cord.

### Optogenetic stimulation and behavior

5 dpf *Tg(gata3:zipACR-YFP)* animals and clutchmates were anesthetized in 0.02% MS-222 and placed on an agar plate under a dissecting microscope. A complete spinal cord transection was made with Vannas spring microscissors, plus a sharpened pin if necessary, between spinal cord segments 2 and 5. Tail blood flow was monitored post-transection and throughout the preparation; animals with significantly reduced blood flow were euthanized and not used for recordings. After transection the animal briefly recovered in extracellular solution and then was embedded in a dorsal up position in 1.2% low melting point agarose. Solidified agarose surrounding the tail caudal to the transection was removed with a dissection scalpel. 200 μM NMDA (Sigma Aldrich) in extracellular solution was added to the dish. Recordings were initiated after tail movement began, typically 2-10 min later.

Behavior experiments were performed with a Scientifica SliceScope upright microscope equipped with a Fastec HiSpec1 camera and an Olympus Plan N 4x/0.10 objective. Image collection was made with Fastec acquisition software. Images were acquired at 200 Hz for 5 seconds. Optical stimulation was made with 100% intensity full field epi-illumination from a CoolLED pe300ultra source routed through a GFP filter cube (Chroma 49002). Recordings with optical stimulation were alternated with recordings without stimulation; n = 6 – 17 recordings for each animal.

Analysis was run in MATLAB R2017a with custom code adapted from Severi et al. [45]. The caudal edge of the transection and the tail periphery were manually selected as tail boundaries and 10 points for tracking were evenly distributed along the body. The caudal-most tail point was used to calculate tail speed (mm/s) at each frame of the recordings. A tail speed threshold of 0.5 mm/s was used to distinguish true movement from tail drift. Tail movement amplitude was calculated as the maximum tail displacement in the initial second of each recording. Tail beat frequency was computed from left-to-right tail oscillations during manually identified movement bouts; 6-30 consecutive peaks were averaged for each recording.

## Acknowledgements

We are grateful to Drs. David McLean and Sandeep Kishore for sharing the *Tg(UAS:Kaede)* and *Tg(gad1b:GFP)* transgenic lines and the UAS:Catch plasmid. Thanks to Dr. Kristen Severi for providing tail tracking code, Marquise Jones for cell counting work, and Dr. Paul Stein for his thoughtful insights. We gratefully acknowledge the Washington University Zebrafish Facility. Imaging experiments were performed in part through the use of Washington University Center for Cellular Imaging (WUCCI) supported by Washington University School of Medicine, The Children’s Discovery Institute of Washington University and St. Louis Children’s Hospital and the Foundation for Barnes-Jewish Hospital. This work was supported by funding through the National Institute of Health (NIH) R00 DC012536 (M.W.B.), R01 DC016413 (M.W.B.), a Sloan Research Fellowship (M.W.B.), The Children’s Discovery Institute of Washington University (M.W.B.), NINDS F32 NS103247 (R.A.C.), and the National BioResource Project in Japan (S.H.). M.W.B. is a Pew Biomedical Scholar and a McKnight Foundation Scholar.

## Supplementary Figures

**Figure S1.**
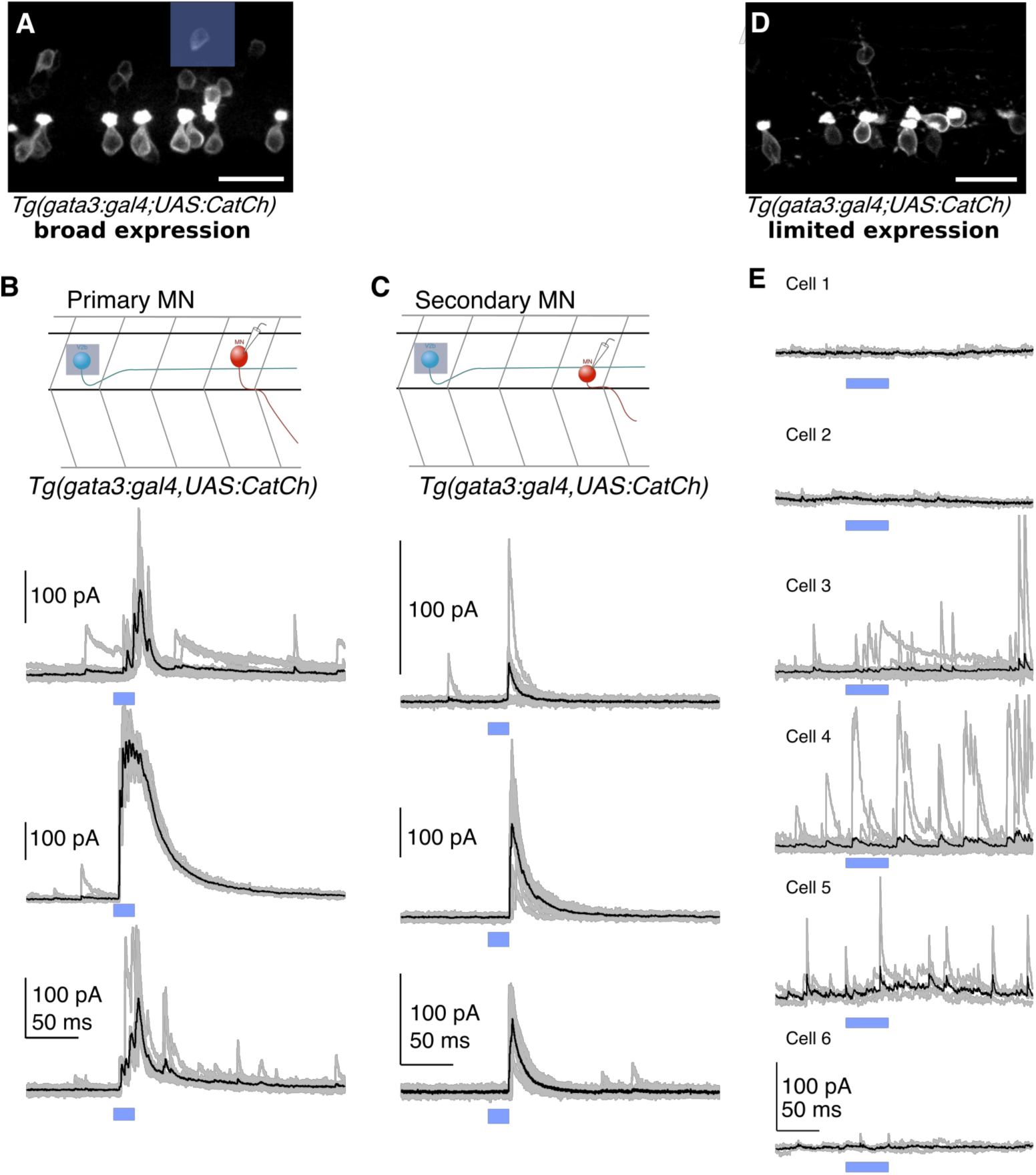
Optogenetically evoked IPSCs originate from V2b neurons, not CSF-cNs. (A) Confocal projection of a hemisegment of spinal cord in a *Tg(gata3:Gal4; UAS:CatCh)* animal with broad CatCh expression. A blue square in the right panel shows an example 20 μm × 20 μm DMD stimulation pattern in which 1-2 V2b cells are targeted. Scale bar = 20 μm. (B) Schematic and whole cell recordings from primary motor neurons in animals with broad CatCh expression, such as in (A). Shorter stimulation times (20 ms) and targeted illumination, as schematized in (A), reliably elicit IPSCs in primary motor neurons. (C) Schematic and recordings from secondary (slow) motor neurons similar to (D). Light stimulation reliably evoked current reponses in secondary motor neurons. (D) Confocal image showing limited CatCh expression in a hemisegment of spinal cord in *Tg(gata3:Gal4; UAS:CatCh)* animals. CatCh is widely expressed in CSF-cN in both animals (A and D), but from animal to animal, there were variations in the intensity of CatCh expression in V2b neurons. Scale bar = 20 μm. (E) Whole cell recordings of primary motor neurons in animals with low V2b CatCh expression. Individual traces are shown in grey and averages in black. Blue bar represents the optical stimulation timing. Longer stimulation times (50 ms) and full field illumination were used to maximize IPSC responses in the recorded cell. Motor neurons receive few light-triggered currents in animals with low CatCh expression in V2b neurons. Together this demonstrates that synaptic currents in Figs. 5 and 6 originate from V2b and not CSF-cN neurons.

**Figure S2.**
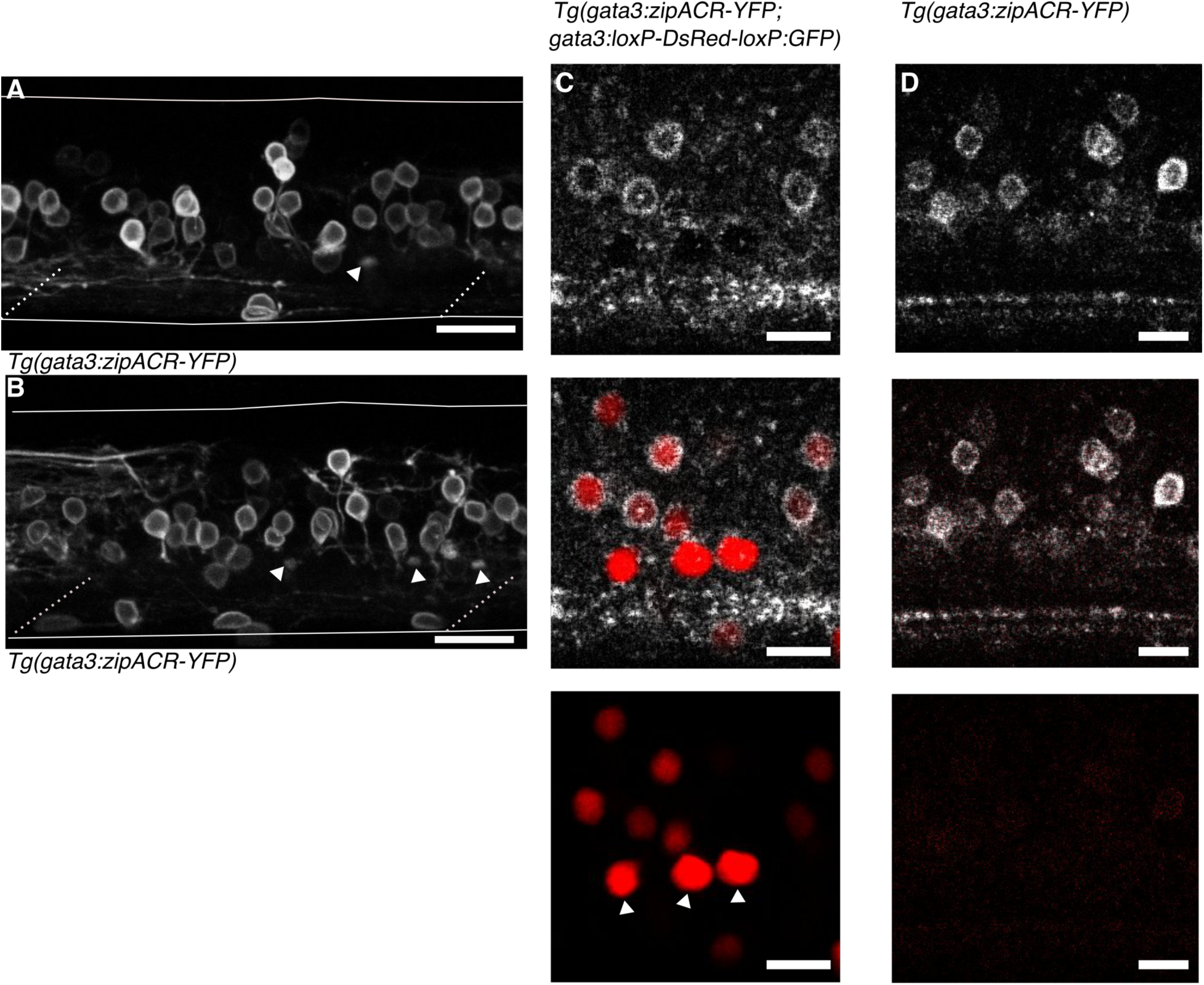
Anatomy of *Tg(gata3:zipACR-YFP)* expression. (A) and (B) depict additional images of Tg(gata3:zipACR-YFP) expression in the full mediolateral extent of the spinal cord in one segment, white triangles mark putative CSF-cN appical extensions into the central canal, noteably CSF-cN soma are not labeled with YFP. Scale bars = 20 μm. (C) Spectrally deconvolved images of *Tg(gata3:zipACR-YFP; gata3:loxP-DsRed-loxP:GFP)*, see Methods. White (top) shows YFP emission and red (bottom) shows DsRed emission. Dorsal CSF-cN are noted with white triangles. CSF-cN somata are distinctly labeled with DsRed (BAC generated line) but not YFP (CRISPR generated line). Scale bar = 10 μm. (D) Example spectral deconvolution of Tg(gata3:zipACR-YFP) showing negligible DsRed emission in control sample. Scale bar = 10 μm.

**Figure S3.**
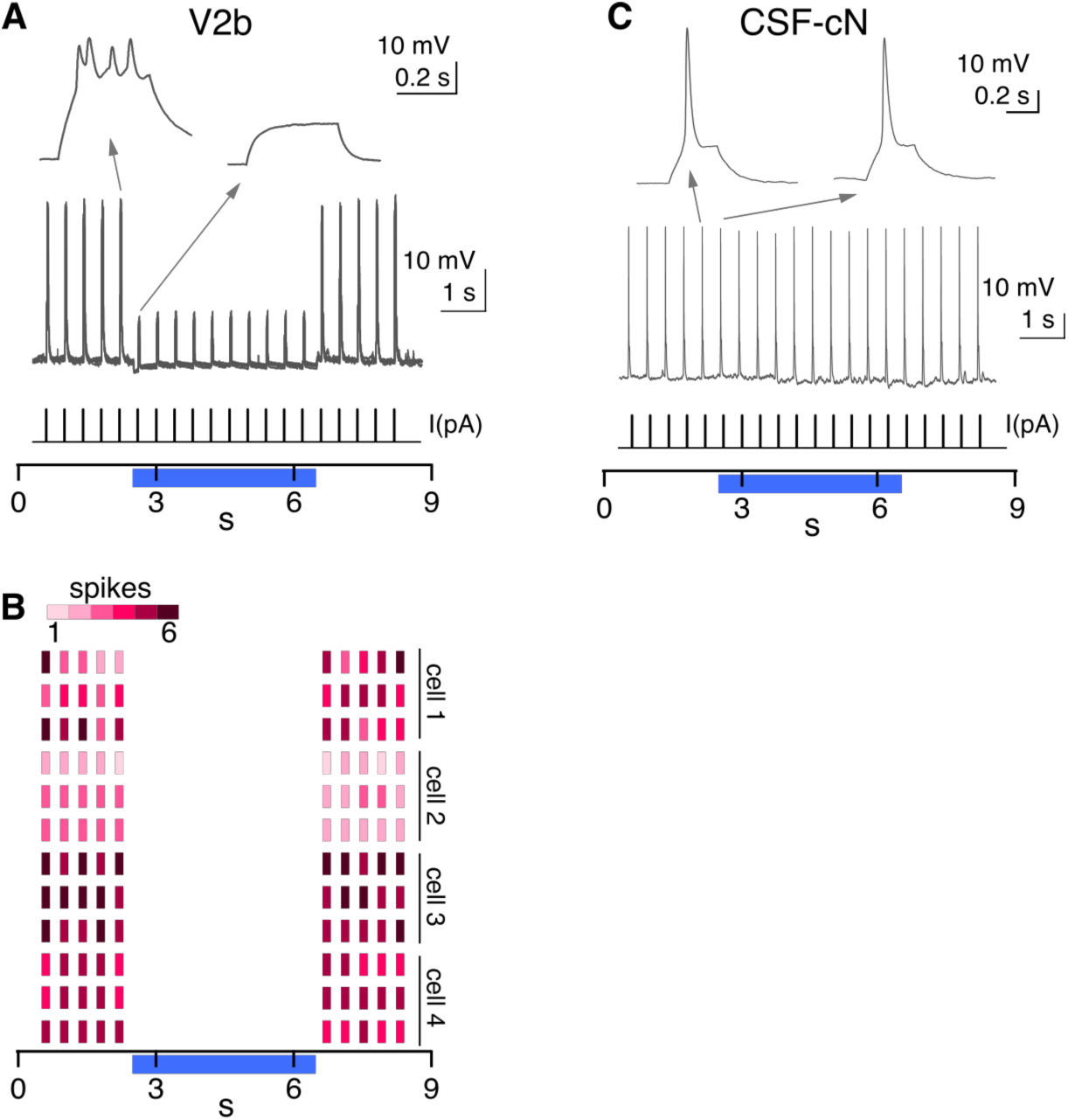
Additional recordings in *Tg(gata3:zipACR-YFP)* animals. (A) A whole cell recording during repeated current steps (20 ms duration) is shown for an example V2b neuron in a Tg(gata3:zipACR-YFP) animal, high intensity light stimulation is provided and indicated by the blue bar. An expanded view of current steps before and during optical stimulation are shown above with arrows. (B) Raster plot of action potentials for 3 trials of 4 V2b cells demonstrates the robust supression of spiking in V2b with high intensity light. Color value represents number of spikes elicited during each current step. (C) A whole cell recording is shown for an example CSF-cN neuron in a Tg(gata3:zipACR-YFP) animal, low intensity light stimulation, similar to Fig. 7, is provided and indicated by the blue bar. An expanded view of current steps before and during optical stimulation are shown above with arrows.

## References

1. Arber, S. (2012). Motor circuits in action: specification, connectivity, and function. Neuron 74, 975–989.

2. Song, J., Dahlberg, E., and El Manira, A. (2018). V2a interneuron diversity tailors spinal circuit organization to control the vigor of locomotor movements. Nat Commun 9, 3370.

3. Menelaou, E., Kishore, S., and McLean, D.L. (2019). Distinct Spinal V2a and V0d Microcircuits Distribute Locomotor Control in Larval Zebrafish. Biorxiv.

4. Menelaou, E., VanDunk, C., and McLean, D.L. (2014). Differences in the morphology of spinal V2a neurons reflect their recruitment order during swimming in larval zebrafish. J Comp Neurol 522, 1232–1248.

5. Hayashi, M., Hinckley, C.A., Driscoll, S.P., Moore, N.J., Levine, A.J., Hilde, K.L., Sharma, K., and Pfaff, S.L. (2018). Graded Arrays of Spinal and Supraspinal V2a Interneuron Subtypes Underlie Forelimb and Hindlimb Motor Control. Neuron 97, 869–884 e865.

6. Bikoff, J.B., Gabitto, M.I., Rivard, A.F., Drobac, E., Machado, T.A., Miri, A., Brenner-Morton, S., Famojure, E., Diaz, C., Alvarez, F.J., et al. (2016). Spinal Inhibitory Interneuron Diversity Delineates Variant Motor Microcircuits. Cell 165, 207–219.

7. Grillner, S., Deliagina, T., El Manira, A., Hill, R.G., Orlovsky, G.N., and Wallén, P. (1995). Neural networks that co-ordinate locomotion and body orientation in lamprey. Trends in Neuroscience 18, 270–279.

8. Danner, S.M., Shevtsova, N.A., Frigon, A., and Rybak, I.A. (2017). Computational modeling of spinal circuits controlling limb coordination and gaits in quadrupeds. Elife 6.

9. Bagnall, M.W., and McLean, D.L. (2014). Modular Organization of Axial Microcircuits in Zebrafish. Science 343, 197–200.

10. Kishore, S., Bagnall, M.W., and McLean, D.L. (2014). Systematic shifts in the balance of excitation and inhibition coordinate the activity of axial motor pools at different speeds of locomotion. J Neurosci 34, 14046–14054.

11. Berg, R.W., Alaburda, A., and Hounsgaard, J. (2007). Balanced inhibition and excitation drive spike activity in spinal half-centers. Science 315, 390–393.

12. Petersen, P.C., Vestergaard, M., Jensen, K.H., and Berg, R.W. (2014). Premotor spinal network with balanced excitation and inhibition during motor patterns has high resilience to structural division. J Neurosci 34, 2774–2784.

13. Li, W.C., and Moult, P.R. (2012). The control of locomotor frequency by excitation and inhibition. J Neurosci 32, 6220–6230.

14. Petersen, P.C., and Berg, R.W. (2016). Lognormal firing rate distribution reveals prominent fluctuation-driven regime in spinal motor networks. Elife 5.

15. Alvarez, F.J., Jonas, P.C., Sapir, T., Hartley, R., Berrocal, M.C., Geiman, E.J., Todd, A.J., and Goulding, M. (2005). Postnatal phenotype and localization of spinal cord V1 derived interneurons. J Comp Neurol 493, 177–192.

16. Sapir, T., Geiman, E.J., Wang, Z., Velasquez, T., Mitsui, S., Yoshihara, Y., Frank, E., Alvarez, F.J., and Goulding, M. (2004). Pax6 and engrailed 1 regulate two distinct aspects of renshaw cell development. J Neurosci 24, 1255–1264.

17. Bhumbra, G.S., Bannatyne, B.A., Watanabe, M., Todd, A.J., Maxwell, D.J., and Beato, M. (2014). The recurrent case for the Renshaw cell. J Neurosci 34, 12919–12932.

18. Britz, O., Zhang, J., Grossmann, K.S., Dyck, J., Kim, J.C., Dymecki, S., Gosgnach, S., and Goulding, M. (2015). A genetically defined asymmetry underlies the inhibitory control of flexor-extensor locomotor movements. Elife 4.

19. Zhang, J., Lanuza, G.M., Britz, O., Wang, Z., Siembab, V.C., Zhang, Y., Velasquez, T., Alvarez, F.J., Frank, E., and Goulding, M. (2014). V1 and v2b interneurons secure the alternating flexor-extensor motor activity mice require for limbed locomotion. Neuron 82, 138–150.

20. McCrea, D.A., and Rybak, I.A. (2008). Organization of mammalian locomotor rhythm and pattern generation. Brain Res Rev 57, 134–146.

21. McLean, D.L., Fan, J., Higashijima, S., Hale, M.E., and Fetcho, J.R. (2007). A topographic map of recruitment in spinal cord. Nature 446, 71–75.

22. Menelaou, E., and McLean, D.L. (2012). A gradient in endogenous rhythmicity and oscillatory drive matches recruitment order in an axial motor pool. J Neurosci 32, 10925–10939.

23. McLean, D.L., Masino, M.A., Koh, I.Y., Lindquist, W.B., and Fetcho, J.R. (2008). Continuous shifts in the active set of spinal interneurons during changes in locomotor speed. Nat Neurosci 11, 1419–1429.

24. Batista, M.F., Jacobstein, J., and Lewis, K.E. (2008). Zebrafish V2 cells develop into excitatory CiD and Notch signalling dependent inhibitory VeLD interneurons. Dev Biol 322, 263–275.

25. Lundfald, L., Restrepo, C.E., Butt, S.J., Peng, C.Y., Droho, S., Endo, T., Zeilhofer, H.U., Sharma, K., and Kiehn, O. (2007). Phenotype of V2-derived interneurons and their relationship to the axon guidance molecule EphA4 in the developing mouse spinal cord. Eur J Neurosci 26, 2989–3002.

26. Kimura, Y., Satou, C., and Higashijima, S. (2008). V2a and V2b neurons are generated by the final divisions of pair-producing progenitors in the zebrafish spinal cord. Development 135, 3001–3005.

27. Eklöf-Ljunggren, E., Haupt, S., Ausborn, J., Dehnisch, I., Uhlén, P., Higashijima, S., and El Manira, A. (2012). Origin of excitation underlying locomotion in the spinal circuit of zebrafish. Proceedings of the National Academy of Sciences 109, 5511–5516.

28. Kimura, Y., Satou, C., Fujioka, S., Shoji, W., Umeda, K., Ishizuka, T., Yawo, H., and Higashijima, S. (2013). Hindbrain V2a neurons in the excitation of spinal locomotor circuits during zebrafish swimming. Curr Biol 23, 843–849.

29. Zhong, G., Sharma, K., and Harris-Warrick, R.M. (2011). Frequency-dependent recruitment of V2a interneurons during fictive locomotion in the mouse spinal cord. Nat Commun 2, 274.

30. Crone, S.A., Zhong, G., Harris-Warrick, R., and Sharma, K. (2009). In mice lacking V2a interneurons, gait depends on speed of locomotion. J Neurosci 29, 7098–7109.

31. Ampatzis, K., Song, J., Ausborn, J., and El Manira, A. (2014). Separate microcircuit modules of distinct v2a interneurons and motoneurons control the speed of locomotion. Neuron 83, 934–943.

32. Karunaratne, A., Hargrave, M., Poh, A., and Yamada, T. (2002). GATA Proteins Identify a Novel Ventral Interneuron Subclass in the Developing Chick Spinal Cord. Developmental Biology 249, 30–43.

33. Wingert, R.A., Selleck, R., Yu, J., Song, H.D., Chen, Z., Song, A., Zhou, Y., Thisse, B., Thisse, C., McMahon, A.P., et al. (2007). The cdx genes and retinoic acid control the positioning and segmentation of the zebrafish pronephros. PLoS Genet 3, 1922–1938.

34. Petracca, Y.L., Sartoretti, M.M., Di Bella, D.J., Marin-Burgin, A., Carcagno, A.L., Schinder, A.F., and Lanuza, G.M. (2016). The late and dual origin of cerebrospinal fluid-contacting neurons in the mouse spinal cord. Development 143, 880–891.

35. Ando, R., Hama, H., Yamamoto-Hino, M., Mizuno, H., and Miyawaki, A. (2002). An optical marker based on the UV-induced green-to-red photoconversion of a fluorescent protein. Proc Natl Acad Sci U S A 99, 12651–12656.

36. Higashijima, S., Mandel, G., and Fetcho, J.R. (2004). Distribution of prospective glutamatergic, glycinergic, and GABAergic neurons in embryonic and larval zebrafish. J Comp Neurol 480, 1–18.

37. Berki, A.C.O.D., M.J.; and Antal, M. (1995). Developmental expression of glycine immunoreactivity and its colocalization with GABA in the embryonic chick lumbosacral spinal cord. Journal of Comparative Neurology 362, 583.

38. Hubbard, J.M., Bohm, U.L., Prendergast, A., Tseng, P.B., Newman, M., Stokes, C., and Wyart, C. (2016). Intraspinal Sensory Neurons Provide Powerful Inhibition to Motor Circuits Ensuring Postural Control during Locomotion. Curr Biol 26, 2841–2853.

39. Chopek, J.W., Nascimento, F., Beato, M., Brownstone, R.M., and Zhang, Y. (2018). Sub-populations of Spinal V3 Interneurons Form Focal Modules of Layered Pre-motor Microcircuits. Cell Rep 25, 146–156 e143.

40. Borowska, J., Jones, C.T., Zhang, H., Blacklaws, J., Goulding, M., and Zhang, Y. (2013). Functional subpopulations of V3 interneurons in the mature mouse spinal cord. J Neurosci 33, 18553–18565.

41. Bergs, A., Schultheis, C., Fischer, E., Tsunoda, S.P., Erbguth, K., Husson, S.J., Govorunova, E., Spudich, J.L., Nagel, G., Gottschalk, A., et al. (2018). Rhodopsin optogenetic toolbox v2.0 for light-sensitive excitation and inhibition in Caenorhabditis elegans. PLoS One 13, e0191802.

42. McDearmid, J.R., and Drapeau, P. (2006). Rhythmic motor activity evoked by NMDA in the spinal zebrafish larva. J Neurophysiol 95, 401–417.

43. Wiggin, T.D., Peck, J.H., and Masino, M.A. (2014). Coordination of fictive motor activity in the larval zebrafish is generated by non-segmental mechanisms. PLoS One 9, e109117.

44. Wiggin, T.D., Anderson, T.M., Eian, J., Peck, J.H., and Masino, M.A. (2012). Episodic swimming in the larval zebrafish is generated by a spatially distributed spinal network with modular functional organization. J Neurophysiol 108, 925–934.

45. Severi, K.E., Böhm, U.L., and Wyart, C. (2018). Investigation of hindbrain activity during active locomotion reveals inhibitory neurons involved in sensorimotor processing. Scientific Reports 8.

46. Bohm, U.L., Prendergast, A., Djenoune, L., Nunes Figueiredo, S., Gomez, J., Stokes, C., Kaiser, S., Suster, M., Kawakami, K., Charpentier, M., et al. (2016). CSF-contacting neurons regulate locomotion by relaying mechanical stimuli to spinal circuits. Nat Commun 7, 10866.

47. McLean, D.L., and Fetcho, J.R. (2009). Spinal interneurons differentiate sequentially from those driving the fastest swimming movements in larval zebrafish to those driving the slowest ones. J Neurosci.

48. Svara, F.N., Kornfeld, J., Denk, W., and Bollmann, J.H. (2018). Volume EM Reconstruction of Spinal Cord Reveals Wiring Specificity in Speed-Related Motor Circuits. Cell Rep 23, 2942–2954.

49. Vergara, H.M., Bertucci, P.Y., Hantz, P., Tosches, M.A., Achim, K., Vopalensky, P., and Arendt, D. (2017). Whole-organism cellular gene-expression atlas reveals conserved cell types in the ventral nerve cord of Platynereis dumerilii. Proc Natl Acad Sci U S A 114, 5878–5885.

50. Francius, C., Harris, A., Rucchin, V., Hendricks, T.J., Stam, F.J., Barber, M., Kurek, D., Grosveld, F.G., Pierani, A., Goulding, M., et al. (2013). Identification of multiple subsets of ventral interneurons and differential distribution along the rostrocaudal axis of the developing spinal cord. PLoS One 8, e70325.

51. Bjornfors, E.R., and El Manira, A. (2016). Functional diversity of excitatory commissural interneurons in adult zebrafish. Elife 5.

52. Hale, M., Ritter, D.A., and Fetcho, J.R. (2001). A Confocal Study of Spinal Interneurons in Living Larval Zebrafish. Journal of Comparative Neurology, 1–16.

53. Bernhardt, R.R., Patel, C.K., Wilson, S.W., and J.Y., K. (1992). Axonal trajectories and distribution of GABAergic spinal neurons in wildtype and mutant zebrafish lacking floor plate cells. Journal of Comparative Neurology 362, 263.

54. Panayi, H., Panayiotou, E., Orford, M., Genethliou, N., Mean, R., Lapathitis, G., Li, S., Xiang, M., Kessaris, N., Richardson, W.D., et al. (2010). Sox1 is required for the specification of a novel p2-derived interneuron subtype in the mouse ventral spinal cord. J Neurosci 30, 12274–12280.

55. Andrzejczuk, L.A., Banerjee, S., England, S.J., Voufo, C., Kamara, K., and Lewis, K.E. (2018). Tal1, Gata2a, and Gata3 Have Distinct Functions in the Development of V2b and Cerebrospinal Fluid-Contacting KA Spinal Neurons. Front Neurosci 12, 170.

56. Gerber, V., Yang, L., Takamiya, M., Ribes, V., Gourain, V., Peravali, R., Stegmaier, J., Mikut, R., Reischl, M., Ferg, M., et al. (2019). The HMG box transcription factors Sox1a and Sox1b specify a new class of glycinergic interneuron in the spinal cord of zebrafish embryos. Development 146.

57. Talpalar, A.E., Bouvier, J., Borgius, L., Fortin, G., Pierani, A., and Kiehn, O. (2013). Dual-mode operation of neuronal networks involved in left-right alternation. Nature 500, 85–88.

58. Li, W.C., Higashijima, S., Parry, D.M., Roberts, A., and Soffe, S.R. (2004). Primitive roles for inhibitory interneurons in developing frog spinal cord. J Neurosci 24, 5840–5848.

59. Ruder, L., Takeoka, A., and Arber, S. (2016). Long-Distance Descending Spinal Neurons Ensure Quadrupedal Locomotor Stability. Neuron.

60. Gosgnach, S., Lanuza, G.M., Butt, S.J., Saueressig, H., Zhang, Y., Velasquez, T., Riethmacher, D., Callaway, E.M., Kiehn, O., and Goulding, M. (2006). V1 spinal neurons regulate the speed of vertebrate locomotor outputs. Nature 440, 215–219.

61. Kohsaka, H., Takasu, E., Morimoto, T., and Nose, A. (2014). A group of segmental premotor interneurons regulates the speed of axial locomotion in Drosophila larvae. Curr Biol 24, 2632–2642.

62. Berg, R.W., Ditlevsen, S., and Hounsgaard, J. (2008). Intense synaptic activity enhances temporal resolution in spinal motoneurons. PLoS One 3, e3218.

63. Pouille, F., Marin-Burgin, A., Adesnik, H., Atallah, B.V., and Scanziani, M. (2009). Input normalization by global feedforward inhibition expands cortical dynamic range. Nat Neurosci 12, 1577–1585.

64. Berg, R.W. (2017). Neuronal Population Activity in Spinal Motor Circuits: Greater Than the Sum of Its Parts. Front Neural Circuits 11, 103.

65. Berkowitz, A. (2002). Both shared and specialized spinal circuitry for scratching and swimming in turtles. J Comp Physiol A Neuroethol Sens Neural Behav Physiol 188, 225–234.

66. Berkowitz, A. (2007). Spinal interneurons that are selectively activated during fictive flexion reflex. J Neurosci 27, 4634–4641.

67. Kimura, Y., Okamura, Y., and Higashijima, S. (2006). alx, a zebrafish homolog of Chx10, marks ipsilateral descending excitatory interneurons that participate in the regulation of spinal locomotor circuits. J Neurosci 26, 5684–5697.

68. Satou, C., Kimura, Y., and Higashijima, S. (2012). Generation of multiple classes of V0 neurons in zebrafish spinal cord: progenitor heterogeneity and temporal control of neuronal diversity. J Neurosci 32, 1771–1783.

69. Kimura, Y., Hisano, Y., Kawahara, A., and Higashijima, S. (2014). Efficient generation of knock-in transgenic zebrafish carrying reporter/driver genes by CRISPR/Cas9-mediated genome engineering. Sci Rep 4, 6545.

70. Kleinlogel, S., Feldbauer, K., Dempski, R.E., Fotis, H., Wood, P.G., Bamann, C., and Bamberg, E. (2011). Ultra light-sensitive and fast neuronal activation with the Ca(2)+-permeable channelrhodopsin CatCh. Nat Neurosci 14, 513–518.

71. Choi, H.M.T., Schwarzkopf, M., Fornace, M.E., Acharya, A., Artavanis, G., Stegmaier, J., Cunha, A., and Pierce, N.A. (2018). Third-generation in situ hybridization chain reaction: multiplexed, quantitative, sensitive, versatile, robust. Development 145.

72. King, R.S., and Newmark, P.A. (2018). Whole-Mount In Situ Hybridization of Planarians. Methods Mol Biol 1774, 379–392.

73. King, R.S., and Newmark, P.A. (2013). In situ hybridization protocol for enhanced detection of gene expression in the planarian Schmidtea mediterranea. BMC Developmental Biology 13.

74. Rothman, J.S., and Silver, R.A. (2018). NeuroMatic: An Integrated Open-Source Software Toolkit for Acquisition, Analysis and Simulation of Electrophysiological Data. Front Neuroinform 12, 14.

75. Schindelin, J., Arganda-Carreras, I., Frise, E., Kaynig, V., Longair, M., Pietzsch, T., Preibisch, S., Rueden, C., Saalfeld, S., Schmid, B., et al. (2012). Fiji: an open-source platform for biological-image analysis. Nat Methods 9, 676–682.

76. Preibisch, S., Saalfeld, S., and Tomancak, P. (2009). Globally optimal stitching of tiled 3D microscopic image acquisitions. Bioinformatics 25, 1463–1465.

77. Longair, M.H., Baker, D.A., and Armstrong, J.D. (2011). Simple Neurite Tracer: open source software for reconstruction, visualization and analysis of neuronal processes. Bioinformatics 27, 2453–2454.

78. Arganda-Carreras, I., Kaynig, V., Rueden, C., Eliceiri, K.W., Schindelin, J., Cardona, A., and Sebastian Seung, H. (2017). Trainable Weka Segmentation: a machine learning tool for microscopy pixel classification. Bioinformatics 33, 2424–2426.

